# BMP6 coordinates dermal papilla and epithelial cell states for hair follicle engineering and regeneration

**DOI:** 10.64898/2026.09.30.755211

**Authors:** Lindsey G. Siegfried, Linli Zhou, Jie Wang, Jennifer M. Hahn, Caroline Dodson, Yeon Sook Choi, Jodie L. Babitt, Dorothy M. Supp, Sarah E. Millar, Thomas Andl, Yuhang Zhang

**Affiliations:** Division of Pharmaceutical Sciences, University of Cincinnati College of Pharmacy, Cincinnati, OH, USA; Department of Surgery, University of Cincinnati College of Medicine, Cincinnati, OH, USA; Dr. Philip Frost Department of Dermatology and Cutaneous Surgery, University of Miami Miller School of Medicine, Miami, FL; Department of Dermatology, University of Pennsylvania, Philadelphia, PA 19104; Nephrology Division and Endocrine Unit, Massachusetts General Hospital, Boston, MA 02114; Shriners Children’s Ohio, Dayton, OH, USA; Department of Stem Cell Biology and Regenerative Medicine, Icahn School of Medicine at Mount Sinai, New York, NY 10029, USA; Department of Oncological Sciences, Icahn School of Medicine at Mount Sinai, New York, NY 10029, USA; Tisch Cancer Institute, Icahn School of Medicine at Mount Sinai, New York, NY 10029, USA; Stravitz-Sanyal Institute for Liver Disease and Metabolic Health, Virginia Commonwealth University, Richmond, VA, USA

**Keywords:** BMP6, dermal papilla, keratinocyte, hair follicle regeneration, epithelial-mesenchymal interaction, hair follicle engineering

## Abstract

Human hair follicle engineering is hindered by the rapid loss of inductive competence in culture-expanded dermal papilla (DP) fibroblasts. Here we show that bone morphogenetic protein 6 (BMP6) maintains and restores DP-associated characteristics while promoting follicular epithelial differentiation by restricting keratinocyte (KC) proliferation and inducing hair follicle-associated marker expression. Human follicle transcriptomics further linked BMP activity to DP identity and follicular epithelial differentiation. Genetic mouse studies demonstrate that BMP6 in Sox2-positive DP cells contributes to normal anagen onset, progression, and maintenance. Gain- and loss-of-function studies in KC–DP organoids show that BMP6 promotes epithelial– mesenchymal organization and spatially patterned follicular structures. *In vivo*, topical BMP6 promotes anagen onset, whereas *BMP6* overexpression restores the hair follicle-generating capacity of late-passage human DP fibroblasts. Together, these findings identify BMP6 as a compartment-dependent regulator that enhances human DP competence and promotes epithelial differentiation, supporting regenerative hair follicle formation and engineering with culture-expanded DP cells.

## Introduction

Human hair follicle regeneration remains a major challenge in regenerative medicine because successful follicle formation requires coordinated interactions between epithelial and mesenchymal cell populations^1,2^. Yet, the signals that coordinate these epithelial–mesenchymal interactions during follicle formation remain incompletely defined. Within this epithelial– mesenchymal unit, dermal papilla (DP) fibroblasts constitute a key inductive mesenchymal population, providing signals that instruct hair follicle morphogenesis and cycling^3^; however, when expanded in culture, DP fibroblasts progressively lose characteristic features, including alkaline phosphatase (AP) activity, versican (VCAN) expression, and their characteristic ability to aggregate, along with a decline in hair-inductive capacity^4,5^. This loss of inductive competence limits their application in human hair follicle engineering^6^. In our organoid model, late-passage DP fibroblasts failed to support the self-organization of mixed DP–keratinocyte (KC) spheroids into polarized, hair peg-like epithelial–mesenchymal structures^7^, indicating loss of epithelial-instructive function during culture expansion. Identifying signals capable of restoring this specialized DP state is therefore a central challenge in hair follicle engineering.

Among the signaling pathways that regulate epithelial–mesenchymal interactions in the hair follicle^8^, bone morphogenetic proteins (BMPs), members of the transforming growth factor-β superfamily, are central regulators of hair follicle morphogenesis, differentiation, and cycling^9^. Within the follicle, BMP activity is spatially and temporally controlled by compartment-specific expression of BMP ligands and receptors, as well as by extracellular antagonists such as noggin, a well-characterized BMP inhibitor^10–16^. This organization enables BMP signaling to exert stage- and lineage-dependent effects, including regulation of stem and progenitor cell activation, differentiation of matrix-derived epithelial lineages, and maintenance of the specialized DP niche^11,14,17,18^. Because BMP signaling regulates both the DP niche and follicular epithelial differentiation, it is possible that a single BMP ligand acting across these compartments could restore DP competence while promoting epithelial organization. However, whether a specific BMP ligand can restore the specialized properties and epithelial-instructive function lost during expansion of human DP fibroblasts remains unclear.

Among BMP ligands, BMP6 is less well characterized than BMP2 and BMP4 in hair follicle biology but represents a particularly compelling candidate because several independent observations place it at the intersection of DP identity and epithelial–mesenchymal communication^19^. Rendl *et al.* showed that recombinant BMP6 preserved selected signature features of cultured murine DP cells and enhanced their hair-inducing activity *in vivo*, while *Bmp6* overexpression maintained DP-associated gene expression and AP activity^11^. Consistent with these findings, we observed that *Bmp6* expression is strongly induced in β-catenin-activated CD133-positive murine DP fibroblasts but suppressed in hair follicles with epithelial *Dkk1* overexpression, linking BMP6 to the Wnt/β-catenin-associated inductive state of the DP. BMP6 also exhibits biochemical properties that distinguish it from other BMP family members, including reduced susceptibility to antagonism by noggin^20–22^, which may favor sustained BMP6 signaling within the hair follicle niche. However, it remains unknown whether BMP6 can restore the epithelial-instructive function lost during expansion of human DP fibroblasts, whether it acts directly on adjacent KCs to promote follicular differentiation, or whether these activities coordinate epithelial–mesenchymal interactions during hair follicle regeneration.

Here, we investigated whether BMP6 can enhance DP inductive competence following its culture-associated attenuation and how BMP6 functions in both the mesenchymal and epithelial compartments of the hair follicle. Using complementary gain- and loss-of-function approaches in human DP fibroblasts and epithelial–mesenchymal organoids, together with KC differentiation studies, conditional mouse studies, and *in vivo* follicle regeneration assays, we examined the role of BMP6 in DP maintenance, follicular epithelial differentiation, hair-cycle progression, and regenerative follicle formation. Our findings identify BMP6 as a regulator that acts in a compartment-specific manner to coordinate DP and epithelial interactions, providing a framework for restoring the regenerative function of culture-expanded DP fibroblasts for human hair follicle engineering.

## Materials and methods

### Transgenic mouse strains

The sex determining region Y-box 2 (*Sox2)-CreER^T2^* mouse strain (B6;129S-*Sox2^tm1(cre/ERT2)Hoch^*/J; Jax: 017593) was obtained from The Jackson Laboratory (Bar Harbor, ME). The *Bmp6^fl/fl^*(Bmp6 floxed) mouse strain was generated in Dr. Jodie Babitt’s laboratory using embryonic stem cells from the Mutant Mouse Resource and Research Center (Lexicon 1E10, MMRRC:029763-UCD)^23^. Both strains were backcrossed onto the C57BL/6 (B6) genetic background. To generate *Sox2-CreER^T2^*; *Bmp6^fl/fl^* mice (DP-*Bmp6**^-/-^***), *Sox2-CreER^T2^* and *Bmp6^fl/fl^*mice were crossed for several generations. Mice were genotyped by polymerase chain reaction (PCR) analysis of genomic DNA extracted from mouse tail biopsies. The *Sox2-CreER^T2^* allele was detected with the following primers: forward, 5′-CAGGAGTTGTCAAGCCAGAGAA-3′; and reverse, 5′-CAGGGTGTTATAAGCAATCCC-3′. *Bmp6* wild-type and floxed alleles were genotyped using the following primers: forward, 5′-TCCATGATCCCTCTAACTCG-3′; and reverse: 5′-GCTGATGACAGCAGCCATTG-3′. All mice were housed in the Laboratory Animal Services facility at the University of Cincinnati under an artificial 12/12 light/dark cycle and allowed free access to food and water. All mouse procedures were approved by the University of Cincinnati Institutional Animal Care and Use Committee under protocol no. 25-08-29-02. Archived cells and hair follicle tissues from the previously described *CD133-CreER^T^*^2^*; Rosa-rtTA; tetO-*Ctnnb1Δ*N* and keratin 5 (*Krt5)-rtTA; tetO-*Dickkopf Wnt signaling pathway inhibitor 1 (*Dkk1)* transgenic mouse studies were used for the BMP6 analyses shown in Fig. 1F and G^24–27^.

**Figure 1.**
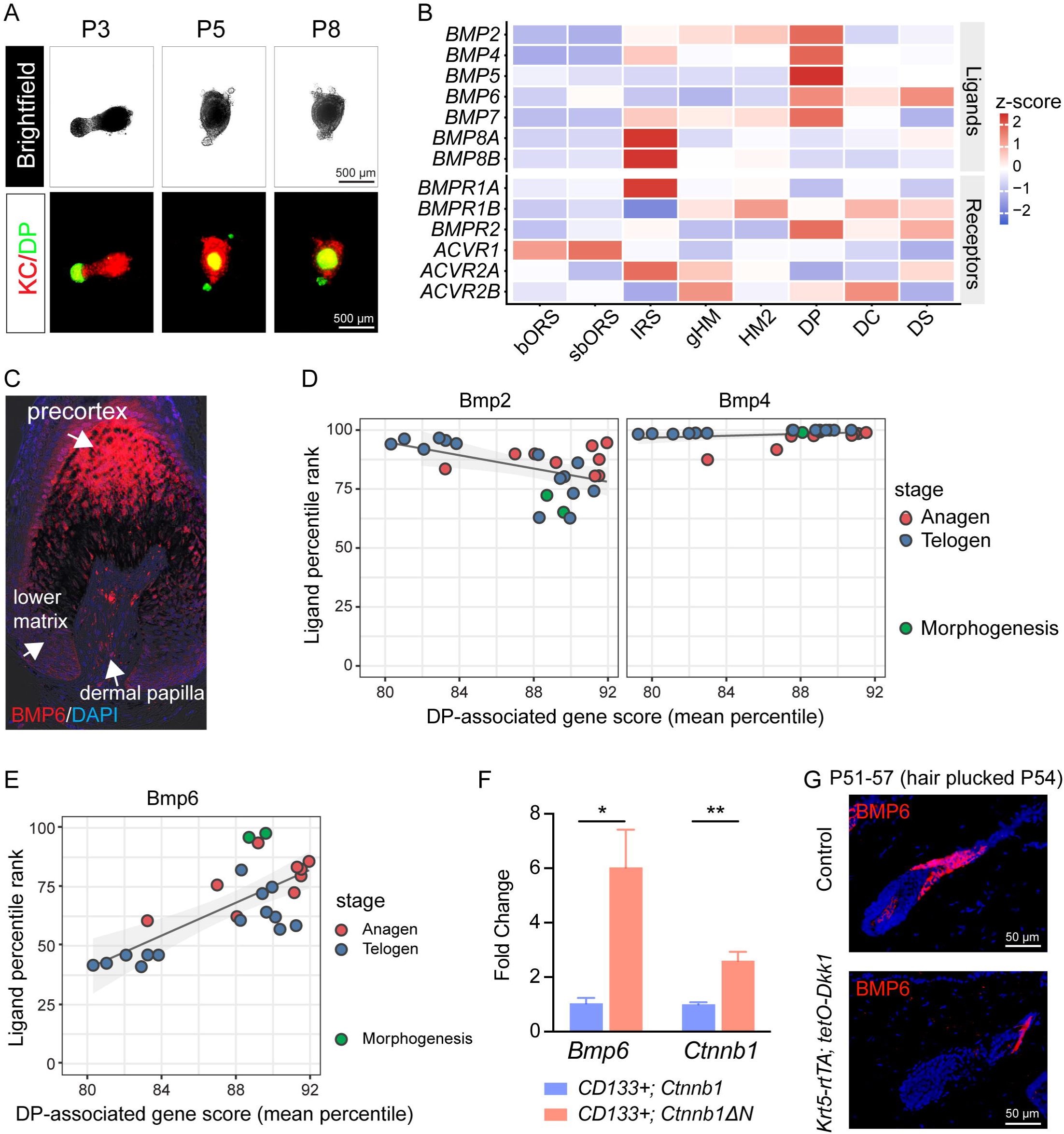
Spatial and molecular evidence associates BMP6 with DP inductive competence. **A.** DP-coordinated self-organization of hair peg-like structures decreases with increasing DP fibroblast passage number in culture. Green fluorescence indicates DP fibroblasts, and red fluorescence indicates KCs. Scale bars = 500 μm. **B.** Human anagen hair follicle transcriptomic data (GSE272333) show predominantly DP-centered expression of *BMP2*, *BMP4*, *BMP5*, and *BMP7*, whereas *BMP6* displays a broader mesenchymal pattern with high relative expression in the DP and DS and lower expression in the DC. **C.** BMP6 immunostaining in human hair follicles shows prominent expression in the precortex in addition to expression in the DP. **D.** Associations of *Bmp2* and *Bmp4* expression with a DP-associated gene score across the murine DP RNA-sequencing datasets. **E.** *Bmp6* expression was positively correlated with the DP-associated gene score across the murine DP RNA-sequencing datasets (R = 0.78, 95% CI 0.55-0.90). **F.** Expression of activated *Ctnnb1*ΔN in CD133-positive murine DP cells significantly increased *Ctnnb1* and *Bmp6* expression by RT-qPCR. Cells were isolated from three pairs of *CD133-CreER^T^*^2^; *Rosa-rtTA; tetO-Ctnnb1*^ΔN^ mice and littermate controls. **G.** Inhibition of epithelial Wnt signaling by Krt5-conditional *Dkk1* overexpression suppresses *Bmp6* expression in murine hair follicles. Scale bar = 50 μm. * *p* < 0.05, ** *p* < 0.01, *** *p* < 0.001, **** *p* < 0.0001; n.s., not significant.

### Induction of Cre expression and conditional *Bmp6* deletion

To induce Cre activity and recombination of *Bmp6* floxed alleles at anagen onset, *Sox2-CreER^T2^*; *Bmp6^fl/fl^* mice and control littermates (*Sox2-CreER^T2^*; *Bmp6^+/+^*, DP-*Bmp6**^+/+^***) were administered 10 mg/mL tamoxifen (Millipore Sigma, #T5648) in corn oil (Thermo Fisher Scientific, #405435000) by intraperitoneal injection at 100 mg/kg body weight from postnatal day 19 (P19) through P26. Skin samples were collected at P35, P38, P40, P42, P45, P50, P55, and P60 as indicated. To ablate *Bmp6* in mid anagen, tamoxifen was administered using the same regimen from P28 through P35. Skin samples were collected at P38, P40, P42, P45, P50, P55, and P60. Mice were euthanized immediately before tissue collection by CO inhalation followed by cervical dislocation as a secondary method. Skin samples were either frozen and embedded in optimal cutting temperature (OCT) compound (Sakura, #4583) or formalin-fixed and paraffin-embedded (FFPE) for histological and immunostaining analyses.

### Hematoxylin and eosin (H&E) staining

FFPE tissues were sectioned at 5 μm. Slides were deparaffinized with xylene substitute (Electron Microscopy Sciences, #64111-04), rehydrated sequentially in 95%, 70%, 50%, and 25% ethanol, followed by water, and stained with hematoxylin (StatLab, SL90-1). Slides were thoroughly washed and treated with differentiation solution (American MasterTech Scientific, #HXD00188E), bluing solution (American MasterTech Scientific, #HXB00242E), and 70% ethanol prior to staining with eosin-phloxine (American MasterTech Scientific, #STE0457). Slides were washed and dehydrated with absolute ethanol, mounted with mounting medium (Vector Laboratories, #H-5000), coverslipped, and imaged on a Nikon Eclipse 80i microscope.

### Murine hair follicle staging

Histologic images from DP-*Bmp6^-/-^* and DP-*Bmp6^+/+^* mice were evaluated for hair follicle stage according to the established murine hair cycle classification system^28^. Follicles were grouped into six categories: early anagen (I–IIIa), mid-anagen (IIIb–IV), late anagen (V–VI)/early catagen (I–III), mid-catagen (IV–VI), late catagen (VII–VIII), and telogen. Five non-overlapping fields were quantified from each of three pairs of DP-*Bmp6^-/-^* and DP-*Bmp6^+/+^*mice at each time point, with individual animals treated as the experimental unit throughout. The anagen-onset conditional knockout cohort was analyzed at P35, P38, P40, P42, P45, and P50, and the mid-anagen conditional knockout cohort at P42, P45, and P50.

To quantify hair cycle progression over time, anagen progression was defined as the proportion of follicles in late anagen/early catagen, mid-catagen, and late catagen, whereas regression was defined as the proportion in mid-catagen and late catagen. Individual biological replicate values were fit by non-linear least-squares regression to a four-parameter logistic model, 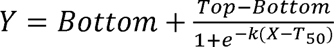 where X represents postnatal day, Bottom and Top were constrained to 0 and 1, respectively, and T_50_ and k were estimated from the data. T_50_ represents the estimated postnatal day at which 50% of follicles reached the defined stage threshold. Genotypes were compared by extra sum-of-squares F test on T_50_. The standard error of ΔT_50_ was calculated as 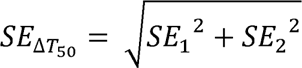 For regression timing, equivalence between genotypes was assessed using a prespecified margin of ± 2 days, based on the temporal resolution of tissue collection, and was concluded when the 95% confidence interval (CI) for ΔT_50_ fell entirely within this margin.

### Primary cell culture

Five human DP fibroblast strains from male and female donors were obtained from PromoCell at passage two (P2) and cultured as previously described in AmnioMAX C-100 medium (Thermo Fisher Scientific, #17001074) supplemented with 10% (v/v) AmnioMAX C-100 supplement (Thermo Fisher Scientific, #12556015) and 1% (v/v) antibiotic/antimycotic (Corning, #30-004-CI) containing 10,000 units/mL penicillin, 10 μg/mL streptomycin, and 25 μg/mL amphotericin B^7^. Medium was replaced every other day, and cells were passaged before reaching 80% confluence using 0.05% trypsin/EDTA (Corning, #25-053-CI). For cell imaging and AP staining in monolayer culture, 5 × 10³ DP fibroblasts were seeded into each well of 8-well chamber slides. All DP fibroblasts used for experimental analyses were between P3 and P8. The donor-derived DP fibroblast strains used in this study and their demographic characteristics are listed in Table S1. Key findings were independently reproduced across multiple donor-derived strains.

Four human female epidermal KC strains were isolated from discarded, de-identified tissue obtained after reduction surgeries in accordance with University of Cincinnati Institutional Review Board-approved protocols, as previously described^29^. Briefly, skin with the subcutaneous fat removed was cut into thin strips and incubated in 2.4 U/mL dispase (Fisher Scientific, #04942078001) in HEPES-buffered saline (HBS) at 4 °C overnight. The following day, the epidermis was peeled from the dermis, washed, and repeatedly agitated in warm 0.025% trypsin (Sigma Aldrich, #T8003)/0.02% EDTA (Sigma Aldrich, #EDS) for two five-minute periods and in HBS for one five-minute period, with released KCs passed through a 70 μm cell strainer after each incubation. Dissociated KCs were pelleted by centrifugation and seeded for culture expansion. KCs were cultured in EpiLife medium (Thermo Fisher Scientific, #MEPI500CA) supplemented with 1% (v/v) EpiLife defined growth supplement (Thermo Fisher Scientific, #S0125) and 1% (v/v) antibiotic/antimycotic. For calcium-induced differentiation experiments, CaCl_2_ was added to the culture media for a final concentration of 2 mM for 48 hours. All KCs used in this study were between P1 and P3. KC identity was confirmed by characteristic morphology and expression of keratin 14 (K14), as previously described^29^. The demographic characteristics of each strain are listed in Table S1.

All cell cultures were maintained in humidified incubators at 37 °C with 5% CO_2_. The phase-contrast images of cultured cells were taken using a Cytation I Cell Imaging Multi-Mode Reader (Bio-Tek). Cell culture reagents were purchased from Thermo Fisher Scientific unless otherwise stated. All experimental procedures involving biosafety hazards were conducted under the University of Cincinnati Institutional Biosafety Committee protocol 25-07-31-01.

### Cell proliferation assay

Cells were seeded into individual wells of flat-bottom 96-well plates (CellTreat, #229196). On days one, three, five, and seven after seeding, cells were stained with 5 μg/mL Hoechst 33342 (Cell Signaling Technology, #4082S) in culture medium for 15 minutes. The staining solution was then removed, and cells were washed three times with PBS. Each well was imaged using a Cytation I Cell Imaging Multi-Mode Reader. Images covering the entire well were acquired, and nuclei were automatically identified and counted based on Hoechst staining using the instrument’s image analysis software. The total number of nuclei per well was used as a measure of cell number at each time point.

### Three-dimensional (3D) DP and KC–DP spheroid culture

3D DP fibroblast-only and KC–DP composite spheroids were prepared as previously described^7^. Briefly, for DP fibroblast-only spheroids, DP fibroblasts at the indicated passages were collected by trypsinization, resuspended in complete DP fibroblast medium, and seeded at 2,500 cells per well in cell-repellent, round-bottom 96- or 384-well microplates (Greiner Bio-One, #650970 and #787979). Spheroids were cultured for three to seven days, with partial medium changes every other day. For KC–DP composite spheroids, DP fibroblasts and KCs were collected by trypsinization and combined in a single-cell suspension containing 2,500 KCs and 2,500 DP fibroblasts in 200 μL of AmnioMAX medium supplemented with 0.2 mM calcium chelator ethylene glycol-bis(2-aminoethylether)-N,N,N′,N′-tetraacetic acid (EGTA; Millipore Sigma, #324626) in addition to 10% AmnioMAX C-100 supplement and 1% antibiotic/antimycotic. The suspension was seeded into each well of cell-repellent, round-bottom 96-well or 384-well plates. Every other day, 100 μL of medium was replaced with fresh composite spheroid medium. Spheroid aggregation and hair peg-like structure formation were assessed regularly using a Cytation I imager as reported^7^.

Fluorescently labeled spheroids were also generated as previously described. CM-DiI (Thermo Fisher Scientific, #D3911) and DiO (Thermo Fisher Scientific, #D275) were prepared in dimethyl sulfoxide (Thermo Fisher Scientific, #BP231) and added to monolayer culture media at 1:1,000 for 30 minutes prior to trypsinization and incorporation into spheroids. Unlabeled controls were performed and no difference in outcome was observed between labeled and non-labeled composite spheroids.

### Recombinant BMP6 treatment

Recombinant BMP6 protein (R&D Systems, #507-BP) was reconstituted in sterile 4 mM HCl with 0.1% bovine serum albumin (BSA; GoldBio, #A-420-100). BMP6 was used at 200 ng/mL for DP fibroblast monoculture, KC–DP composite spheroids, and for seven-day KC monoculture. BMP6 was used at 500 ng/mL for three-day KC monoculture. Cells and spheroids were treated with BMP6 for three to seven days, as indicated. All BMP6-treated DP fibroblasts were at P3 or P5 and all BMP6-treated KCs were between P1 and P3.

### Generation of BMP6-CRISPRa DP fibroblasts

BMP6 single-guide RNAs (sgRNAs) and a non-targeting control sgRNA were designed using the Broad Institute GPP sgRNA Designer^30,31^. The selected sgRNAs targeted a compact, GC-rich region immediately upstream of the BMP6 reference transcription start site, located at chr6:7,726,099 in the GRCh38 genome assembly^32^. The BMP6 and control sgRNA sequences can be found in Table S2. All three BMP6 sgRNAs and the non-targeting control sgRNA were synthesized and cloned in pGenlenti lentiviral vectors carrying puromycin resistance and ZsGreen fluorescence by Genscript (Piscataway, NJ). The dCas9-VPR construct, pLV[Exp]-EFS>dCas9/VPR(ns):P2A:Bsd, was designed and cloned by VectorBuilder (Chicago, IL, USA).

To generate BMP6-CRISPRa and non-targeting control (DPF-sgCtl) DP fibroblast strains, 1 × 10^5^ DP fibroblasts at P2 were first transduced with 50 μL of viral suspension mixed with 450 μL of basal DMEM supplemented with 10 μg/mL polybrene (Santa Cruz Biotechnology, #sc-134220). After 16 hours, cells were washed with sterile PBS, and the transduction medium was replaced with fresh complete AmnioMAX medium. Transduced cells were selected with 10 μg/mL blasticidin (Santa Cruz Biotechnology, #sc-495389) for three days. Blasticidin-resistant cells were subsequently transduced with lentivirus carrying one of the three BMP6 sgRNAs or the non-targeting control sgRNA following the same procedure. ZsGreen expression was visualized on a Cytation I imager. Cells were then selected with 10 μg/mL puromycin (Thermo Fisher Scientific, #A11138-03) for three days. BMP6 induction by the three sgRNAs was compared by Western blotting. The sgRNA that achieved the highest level of BMP6 activation, sg*BMP6*-3, was used for all experiments and designated DPF-sg*BMP6*. All DPF-sg*BMP6* and DPF-sgCtl used for experiments were at P5.

### Generation of direct BMP6-overexpressing DP fibroblasts

The BMP6-overexpressing lentiviral vector, pLV[Exp]-Puro-EF1A>hBMP6:P2A:ZsGreen, and the corresponding empty control vector, pLV[Exp]-Puro-EF1A>ZsGreen1, were also designed and cloned by VectorBuilder. Both plasmids were packaged into third-generation lentiviral particles by the Viral Vector Core Facility at Cincinnati Children’s Hospital Medical Center.

To generate BMP6-overexpressing and control DP fibroblasts, 1 × 10^5^ DP fibroblasts at P3 were seeded in 10 cm dishes, transduced with the corresponding lentiviruses following the same procedure described above, and selected with 10 μg/mL puromycin for three days. The resulting cell lines were designated DPF-Lv*BMP6* and DPF-LvCtl, respectively. BMP6 overexpression was validated by Western blotting, RT-qPCR, or both. All DPF-Lv*BMP6* and DPF-LvCtl cells used for experiments were at P6**–**P7.

### Generation of inducible BMP6-deficient DP fibroblasts

To silence BMP6 expression, human DP fibroblasts were transduced with lentiviral particles carrying doxycycline-inducible *BMP6*-targeting shRNAs (Horizon Discovery, V3SH7669-229470456, V3SH7669-226416669, and V3SH7669-225337503). A non-targeting control shRNA lentivirus was used to generate control DP fibroblasts. Expression of the *BMP6*-targeting or non-targeting shRNA was induced by doxycycline treatment as indicated.

Briefly, P3 DP fibroblasts (3 × 10^5^) were seeded in 10 cm tissue culture dishes and transduced with 25 µL of lentiviral particles with 10 μg/mL polybrene for 16 hours. The virus-containing medium was then replaced with fresh complete AmnioMAX medium. Transduced cells were selected with 10 μg/mL puromycin for three days and subsequently treated with 500 ng/mL doxycycline (Fisher Scientific, #BP2653-1) for 72 hours to induce shRNA and GFP expression. GFP fluorescence was used to confirm induction of the lentiviral construct, and BMP6 knockdown efficiency was evaluated by Western blotting. Among the three shRNAs tested, V3SH7669-225337503 produced the greatest BMP6 knockdown and was selected for all subsequent experiments. Cells carrying V3SH7669-225337503 were designated DPF-sh*BMP6*. DPF-sh*BMP6* and non-targeting control (DPF-shCtl) DP fibroblasts were used at early passage (before P5) to examine the effects of BMP6 depletion while DP-associated characteristics were still relatively retained following transduction at P3.

### Hydrogel culture

To assess clonal expansion and AP positivity in hydrogel culture, 1 × 10^4^ cultured DP fibroblasts were collected by trypsinization, mixed with 100 μL of Extracel hydrogel (Advanced BioMatrix) according to the manufacturer’s instructions, and seeded in each well of a flat-bottomed 96-well plate (CellTreat, #229196). Cultures were maintained in a humidified incubator at 37 °C with 5% CO until the indicated time points. For colony-size quantification, the area of each colony larger than 100 μm² in randomly selected bright-field images was measured using ImageJ (NIH). The mean colony area for each field was calculated and used for comparisons among experimental groups.

### Reverse Transcription Quantitative Real-Time Polymerase Chain Reaction (RT-qPCR)

Cells cultured in 60 mm dishes were harvested by trypsinization. Total RNA was isolated using the RNeasy Plus Micro Kit (Qiagen, #74034) or RNeasy Plus Mini Kit (Qiagen, #74134) for samples containing fewer than 5 × 10^5^ cells or at least 5 × 10^5^ cells, respectively. RNA concentration and purity were assessed using a NanoDrop spectrophotometer (Thermo Fisher Scientific). cDNA was synthesized using the QuantiTect Reverse Transcription Kit (Qiagen, #205311). qPCR was performed using PowerUp SYBR Green Master Mix (Thermo Fisher Scientific, #A25742) on a QuantStudio Real-Time PCR System (Thermo Fisher Scientific). The thermal cycling conditions consisted of incubation at 50 °C for two minutes and polymerase activation at 95 °C for two minutes, followed by 40 cycles of denaturation at 95 °C for 15 seconds and annealing/extension at 60 °C for one minute. Primer pairs for all genes are listed in Table S3. Relative gene expression and fold change were normalized to *ACTB* (DP fibroblasts), *Actb* (murine DP cells), or *GAPDH* (KCs) and calculated using the 2^−ΔΔCt^ method. Fold change in paired samples was calculated against each respective control using the 2^-ΔΔCt^ method.

### Western blotting

Cell pellets were lysed in RIPA buffer (Thermo Fisher Scientific, #89901) supplemented with 1% (v/v) protease and phosphatase inhibitor cocktail (Thermo Fisher Scientific, #78443), with vigorous vortexing between three 10-minute incubations on ice. Lysates were centrifuged at 15,000 × *g* for 15 minutes at 4 °C, and supernatants were collected. Protein concentrations were determined using a bicinchoninic acid (BCA) assay (Thermo Fisher Scientific, #A55860). Proteins (20 μg per lane) were separated by SDS-PAGE on freshly prepared 10% polyacrylamide gels and transferred to nitrocellulose or methanol-activated PVDF membranes using a Trans-Blot Turbo system (Bio-Rad). Membranes were blocked for one hour at room temperature with 5% nonfat dry milk or 5% bovine serum albumin (BSA) in Tris-buffered saline containing 0.5% Tween 20 (TBST; Amresco, #0777-1L), followed by incubation with primary antibodies in blocking buffer overnight at 4 °C.

Primary antibodies included BMP6 (R&D Systems, #MAB507, 1:500), cyclin A2 (Cell Signaling Technology, #4656S, 1:1,000), cyclin B1 (Cell Signaling Technology, #12231S, 1:1,000), cyclin D3 (Cell Signaling Technology, #2936P, 1:1,000), cyclin E (Thermo Fisher Scientific, #MA5-14336, 1:1,000), CDK4 (Cell Signaling Technology, #12790P, 1:1,000), CDK2 (Cell Signaling Technology, #2546P, 1:1,000), actin (Abcam, #ab3280, 1:1,000), and tubulin (Thermo Fisher Scientific, #MA5-16308, 1:5,000). Membranes were washed with TBST and incubated for one hour at room temperature with IRDye 800CW donkey anti-rabbit IgG (LI-COR, #926-32213) or IRDye 680RD donkey anti-mouse IgG (LI-COR, #926-68072) secondary antibodies diluted 1:2,000 in blocking buffer. After washing with TBST, membranes were imaged using an Odyssey CLx imaging system (LI-COR, Nebraska, US).

### AP staining

Cells, KC–DP spheroids, and hydrogel cultures were fixed with 4% paraformaldehyde (PFA; Thermo Fisher Scientific, #J61899-AK) and washed thoroughly with PBS. Samples were then rinsed with 0.1 M Tris-HCl (pH 8.8). Vector Blue or Vector Red AP substrate reagents (Vector Laboratories, #SK-5300 and #SK-5100) were prepared in the same buffer according to the manufacturer’s instructions by combining two drops of each kit reagent in 5 mL of 0.1 M Tris-HCl (pH 8.8). Samples were incubated in staining solution for one hour, washed once with Tris-HCl and twice with PBS, and imaged. Spheroids and organoids were whole-mount imaged in PBS on a Leica DMi8 widefield fluorescence microscope, slide-mounted cells and tissues were imaged on a Nikon Eclipse 80i microscope, and hydrogel cultures were imaged on a Cytation I imager.

For AP quantification in monolayer culture, bright-field images were randomly acquired from each well and analyzed using ImageJ (NIH). For each image, the total cell-covered area and the area positive for blue AP staining were quantified. For exogenous BMP6 treatment, AP activity was expressed as the total AP-positive area. For experiments involving endogenous *BMP6* modulation, AP positivity was calculated as the percentage of AP-positive area relative to the total cell-covered area.

For AP quantification in hydrogel cultures, paired brightfield and fluorescence images were acquired from randomly selected fields. Images were analyzed using ImageJ to determine the total number of colonies in bright-field images and the number of AP-positive colonies in the corresponding red-fluorescence images. AP positivity was calculated as the percentage of AP-positive colonies relative to the total number of colonies in each field.

### Immunostaining

Monolayer cells (5,000–10,000 cells/well) cultured in 8-well chamber slides were fixed in 4% PFA for 15 minutes, washed with PBS, and permeabilized with 0.1% Triton X-100 (Thermo Fisher Scientific, #85111) in PBS for 10 minutes. Spheroids were fixed with 4% PFA and sequentially cryoprotected in 15% and 30% sucrose for one hour each before embedding in OCT compound. Mouse skin tissues were either fixed with 4% PFA and embedded in OCT or FFPE-processed. All sectioning was performed at 5 μm thickness.

Frozen sections were fixed in ice-cold methanol (Thermo Fisher Scientific, #176840025) for eight minutes followed by acetone (Fisher Scientific, #A18-4) for three minutes and washed with PBS to remove OCT. FFPE slides were deparaffinized and sequentially rehydrated following the previously described procedure. Antigen retrieval was performed in 10 mM sodium citrate buffer (pH 6.0) containing 0.5% Tween 20 at 95 °C in a water bath or by microwave heating. To reduce melanin interference, tissue sections were bleached for 45 minutes under bright light in PBS containing 4.5% H O (Fisher Scientific, #BP2633-500) and 2.5% 1 M NaOH. Samples were blocked with 10% BSA in PBS and incubated overnight with primary antibodies diluted in blocking solution. Primary antibodies included hair cortex cytokeratin (AE13; Santa Cruz Biotechnology, #sc-57012, 1:50), BMP6 (Santa Cruz Biotechnology, #sc-7406, 1:50), GATA-binding factor 3 (GATA3; Santa Cruz Biotechnology, #sc-268, 1:50), K6 (Invitrogen, #MA5-32318, 1:1,000), K10 (Novus Biologicals, #NBP2-47650, 1:100), Ki67 (BD Pharmingen, #550609, 1:50), lymphoid enhancer-binding factor 1 (LEF1; Abcam, #ab137872, 1:100), SOX2 (Santa Cruz Biotechnology, #sc-365823, 1:50), trichohyalin (TCHH; gift from Dr. Tung-Tien Sun, NYU Langone Medical Center, 1:50), and VCAN (Invitrogen, #PA1-1748A (human samples); Millipore Sigma, #AB1033; 1:100 (mouse samples)).

For immunofluorescence staining, {Calvo, 2013 #10}secondary antibodies, including anti-rabbit DyLight 594 (Vector Laboratories, #DI-1094), anti-rabbit Alexa Fluor 488 (Invitrogen, #A11034), anti-mouse Alexa Fluor 594 (Invitrogen, #A11005), and anti-mouse Alexa Fluor 488 (Invitrogen, #A32723), were diluted 1:500 in BSA blocking solution. Sections were incubated with the diluted secondary antibody solution for one hour at room temperature, mounted with DAPI-containing mounting medium (Abcam, #ab104139), and imaged on a Nikon Eclipse 80i microscope. For organoid cryosections, nuclei were counterstained with 10 mg/mL Hoechst 33342 diluted in the secondary antibody solution at 1:500. To quantify Ki67-positive cells in immunofluorescence images, the numbers of Ki67-positive nuclei and total DAPI-positive nuclei were counted in each field using ImageJ (NIH). The percentage of Ki67-positive cells was calculated as the number of Ki67-positive nuclei divided by the total number of DAPI-positive nuclei × 100%. For quantification of multinucleated KCs, cells cultured on chamber slides were incubated with rhodamine phalloidin (Invitrogen, #R415; 1:500) and Hoechst 33342 (1:500) to visualize F-actin and nuclei, respectively, allowing identification of individual cell borders and multinucleated cells. Nuclei/actin images were quantified by manually counting the total number of cells and the number of cells containing multiple nuclei.

### Topical BMP6 treatment

Dorsal hair of C57BL/6 mice was shaved at P21 for females and P19 for males. A total of six male and six female mice were used. Two 1 × 2 cm treatment areas were marked on opposite sides of the dorsal skin. The recombinant BMP6 solution was prepared as described and diluted in sterile water. One area of dorsal skin received recombinant BMP6 (1 μg/cm²), whereas the contralateral area received sterile water alone as a vehicle control. For each treatment, 20 μL of solution was evenly applied to the marked skin surface using a 200 μL pipette tip. Treatments were administered once daily for 10 consecutive days, during which mice were monitored for hair follicle anagen entry. Skin biopsies were collected after completion of the 10-day treatment for histological and immunostaining analyses. Hair bulb diameter was quantified in three of the twelve treated mice. Histological hair bulb diameter was quantified in ImageJ by drawing a line across the maximal width of each hair bulb within each field of view and determining the line length using the Measure function. Images were analyzed without spatial calibration; therefore, measurements are reported in arbitrary units (a.u.).

### Patch assay

For the patch assay, trunk skin from P2 C57BL/6 neonates was collected and separated into epidermal and dermal fractions. Skin was floated epidermis-side up in 0.25% trypsin at 37 °C for one hour to separate the epidermis from the dermis. The epidermis was minced and dissociated into a single-cell suspension by vigorous pipetting. The dermis was minced and digested in 0.25% collagenase I (Gibco, #17100-017) and 0.25% collagenase IV (Gibco, #17104-019) for one hour at 37 °C, with manual disruption using a serological pipette every 15 min. Cell suspensions were sequentially filtered through 70-μm and 40-μm strainers, washed with PBS containing 10% FBS and 1% antibiotic/antimycotic, and resuspended in 1 mL PBS containing 2% FBS and 1% antibiotic/antimycotic for cell counting.

P7 human DP fibroblasts overexpressing BMP6 (DPF-Lv*BMP6*) and vector-transduced control (DPF-LvCtl) DP fibroblasts were collected by trypsinization. For each patch, 200 μL of mixed cell suspension containing 1 × 10^5^ DPF-Lv*BMP6* or DPF-LvCtl DP fibroblasts, 1 × 10^6^ neonatal murine epidermal cells, and 5 × 10^5^ neonatal murine dermal cells was injected intradermally into the dorsal skin of nude mice. For positive-control patches, 200 μL of mixed cell suspension containing 1 × 10^5^ P3 human DP fibroblasts, 1 × 10^6^ neonatal murine epidermal cells, and 5 × 10^5^ neonatal murine dermal cells was injected. For negative-control patches, 200 μL containing 1 × 10^6^ neonatal murine epidermal cells and 5 × 10^5^ neonatal murine dermal cells without human DP fibroblasts was injected. For the DPF-Lv*BMP6* versus DPF-LvCtl comparison, eight mice received DPF-Lv*BMP6* on one side and DPF-LvCtl on the contralateral side. For quantitative hair-count analysis, three of the eight paired mice were analyzed. Six P3 DP-positive-control patches and five mouse-cell-only negative-control patches were included. Patches were collected 28 days after injection for assessment of *de novo* hair follicle formation. Both male and female mice were used.

### Murine RNA-Seq data analysis

Processed expression data from four murine DP RNA-seq datasets (GSE109256, GSE300176, GSE235480, GSE136996) were obtained from Gene Expression Omnibus (GEO). Only wild-type or control samples from freshly isolated or microdissected DP were included. Gene identifiers were harmonized to gene symbols using the org.Mm.eg.db 3.20.0 annotation package^33^. To place datasets on a common scale without assuming comparable absolute values, expression was converted to percentile rank within each sample by ranking the normalized expression of each gene relative to all genes in that sample and dividing by the total number of genes. Due to within-sample ranking, transcript length effects are constant per gene and do not contribute to the reported associations.

A DP-associated gene score was calculated as the mean percentile rank of 17 representative DP-associated genes (*Alpl, Sox2, Sox18, Crabp1, Vcan, Lef1, Wnt5a, Fgf7, Fgf10, Igf1, Hgf, Foxd1, Prdm1, Ltbp1, Rspo3, Wif1,* and *Sostdc1*). All 17 genes were present in all datasets. Associations between *Bmp2, Bmp4,* and *Bmp6* expression and the DP-associated gene score were assessed across *n* = 25 samples spanning seven postnatal timepoints in the four datasets using Pearson correlation, because the percentile ranks were approximately continuous and the relationships were approximately linear. Correlation coefficients are reported with 95% confidence intervals. Analyses were performed in R version 4.4.1^34^ using GEOquery 2.74.0^35^, AnnotationDbi 1.68.0^36^, tidyr 1.3.2^37^, ggplot2 4.0.3^38^, and dplyr 1.2.1^39^.

### Human RNA-Seq data analysis

Compartment-resolved RNA-sequencing data from laser-capture microdissected human anagen hair follicles (GSE272333) were obtained from Gene Expression Omnibus (GEO)^40^. This dataset included 3–5 hair follicles each from 7–10 male and female donors, for a total of 110 samples. Compartments included the bulge outer root sheath (bORS), suprabulge outer root sheath (sbORS), inner root sheath (IRS), germinative hair matrix (gHM), precortex matrix (HM2), dermal papilla (DP), lower dermal cup (DC), and dermal sheath (DS). Differential expression was analyzed using DESeq2 with donor included as a blocking factor (∼ donor + compartment). Genes with ≥ 10 counts in ≥ 5 samples were retained, and variance-stabilizing transformation (VST) was applied for downstream analyses. Pairwise compartment comparisons were performed using DESeq2 with log_2_ fold-change shrinkage, with adjusted *p* < 0.05 and |log_2_FC| > 1 used to define differential expression. For compartment-level analyses, mean VST expression was calculated for each gene within each compartment. The BMP ligand and receptor heatmap was displayed as gene-wise z-scores and included *BMP2, BMP4, BMP5, BMP6, BMP7, BMP8A, BMP8B, BMPR1A, BMPR1B, BMPR2, ACVR1, ACVR2A,* and *ACVR2B*. The gHM and HM2 cell-cycle heatmap displayed mean VST expression for *CDKN1A, CDKN2A, CDK2, CDK4, CDK6, CCNA1, CCNB2, CCND3, CCNE1,* and *CCNE2*. Heatmaps were generated using ComplexHeatmap.

A BMP pathway score was calculated for each sample as the mean VST expression of eight BMP-responsive genes (*ID1, ID2, ID3, ID4, SMAD6, SMAD7, BAMBI,* and *JUNB*). Associations between the BMP pathway score and genes of interest were assessed across gHM and HM2 samples combined using Spearman correlation, as VST values are not assumed to have a linear relationship. Mediation analysis was used to examine associations among the BMP pathway score, *LEF1*, and *KRT31*, using *LEF1* ∼ BMP pathway score + donor as the mediator model and *KRT31* ∼ BMP pathway score + *LEF1* + donor as the outcome model. Analyses were performed across all samples and within gHM and HM2, with 1,000 bootstrap simulations used to estimate indirect, direct, and total effects with 95% confidence intervals. These analyses describe relationships within the transcriptomic dataset and do not establish causality. Exploratory network analysis was performed using GENIE3 on VST-normalized expression of selected BMP-responsive and follicular differentiation-associated genes together with the BMP pathway score. The top 20 relationships ranked by GENIE3 weight were retained for visualization. Analyses were performed in R version 4.4.1^34^ using GEOquery 2.74.0^35^, DESeq2 1.46.0^41^, AnnotationDbi 1.68.0^36^, org.Hs.eg.db 3.20.0^42^, tidyr 1.3.2^37^, ggplot2 4.0.3^38^, ComplexHeatmap 2.22.0^43^, mediation 4.5.1^44^, GENIE3 1.28.0^45^, and dplyr 1.2.1^39^.

### Statistical analysis

Data were analyzed using GraphPad Prism 10 (GraphPad Software Inc.) and are presented as mean ± SEM. Comparisons between two groups were performed using unpaired Student’s *t* tests. For the topical BMP6 hair bulb analysis, individual hair follicle measurements from vehicle- and BMP6-treated sites were pooled by treatment condition across three mice and compared using an unpaired Student’s *t* test; the total numbers of follicles analyzed are reported in Fig. 8D. For the patch assay, a within-mouse log fold change was calculated by normalizing each BMP6-overexpressing patch to the contralateral control patch and assessed by a one-sample *t* test. Comparisons between more than two groups were performed using one-way analysis of variance (ANOVA), followed by Dunnett’s multiple comparisons test. The statistical test, sample size, and experimental unit for each analysis are specified in the methods, figures, or corresponding figure legends.

## Results

### Spatial and molecular evidence associates BMP6 with DP inductive competence

We previously showed that composite spheroids generated from P3 human DP fibroblasts and epidermal KCs self-organize into hair peg-like structures containing a proximal region of intermingled DP fibroblasts and KCs and an elongated KC-rich protrusion^7^. This self-organizing capacity declined with DP fibroblast passage and was largely lost beyond P3, consistent with culture-associated attenuation of the DP phenotype (Fig. 1A). Because hair follicle engineering requires expansion of DP fibroblasts while retaining their hair-inductive competence, we sought factors capable of restoring this competence in late-passage cells.

Previous studies in mouse hair follicles have implicated BMP6 in maintaining DP characteristics and regulating follicular epithelial behavior^11,46^. We therefore examined the spatial distribution of BMP ligands and receptors in human anagen follicles using compartment-resolved RNA-seq data (GSE272333), generated from laser-capture microdissected human anagen VI follicles^40^. Whereas BMP2, BMP4, BMP5, and BMP7 showed predominantly DP-centered expression, BMP6 displayed a broader mesenchymal distribution, with high relative expression in both the DP and DS and lower expression in the DC (Fig. 1B). Meanwhile, BMP receptor components were broadly distributed across epithelial and mesenchymal follicular compartments. Immunostaining further confirmed BMP6 expression in the DP and prominent expression in the precortex of human anagen hair follicles (Fig. 1C).

Complementing the human spatial analysis, analysis of four murine DP RNA-sequencing datasets spanning anagen, telogen and morphogenesis showed that comparable associations were not observed for *Bmp2* or *Bmp4* (Fig. 1D), whereas *Bmp6* expression was positively correlated with a DP-associated gene score (Fig. 1E; R = 0.78, 95% CI 0.55–0.90, *p* < 0.0001). In addition, *Bmp6* was strongly increased in CD133-positive DP fibroblasts expressing activated ΔN-b-catenin but suppressed in mouse hair follicles expressing the Wnt inhibitor *Dkk1* (Fig. 1F, G)^24,26^. Together, these findings associate BMP6 with DP-associated molecular characteristics and support further investigation of its role in DP–epithelial interactions. We therefore first asked whether DP-derived BMP6 is required for coordinated follicular epithelial differentiation and hair-cycle progression *in vivo*.

### DP-derived BMP6 is required for sustained anagen and follicular differentiation

Sox2 is an established marker of DP cells in several major mouse hair follicle types, including guard, awl, and auchene follicles^46,47^. To determine whether DP-derived Bmp6 is required for follicular cycling *in vivo*, we generated *Sox2-CreER^T^*^2^; *Bmp6*^fl/fl^ mice and induced *Bmp6* deletion at P19, near entry into the synchronized postnatal anagen (Fig. 2A). Control follicles progressively developed enlarged, deeply positioned, pigmented bulbs characteristic of mature anagen, whereas DP-*Bmp6^-/-^*follicles entered anagen but showed markedly delayed progression and rarely attained comparable late-anagen morphology (Fig. 2B). Quantitative staging confirmed a substantial delay in progression to late anagen/early catagen in mutant follicles (Fig. 2C, D). In contrast, progression into catagen occurred on a similar timetable in control and mutant follicles (Fig. 2E), suggesting that follicles in DP-*Bmp6^-/-^* mice entered regression before completing full anagen maturation. These findings indicate that Bmp6 in Sox2-expressing DP cells contributes to hair follicle anagen onset and progression.

**Figure 2.**
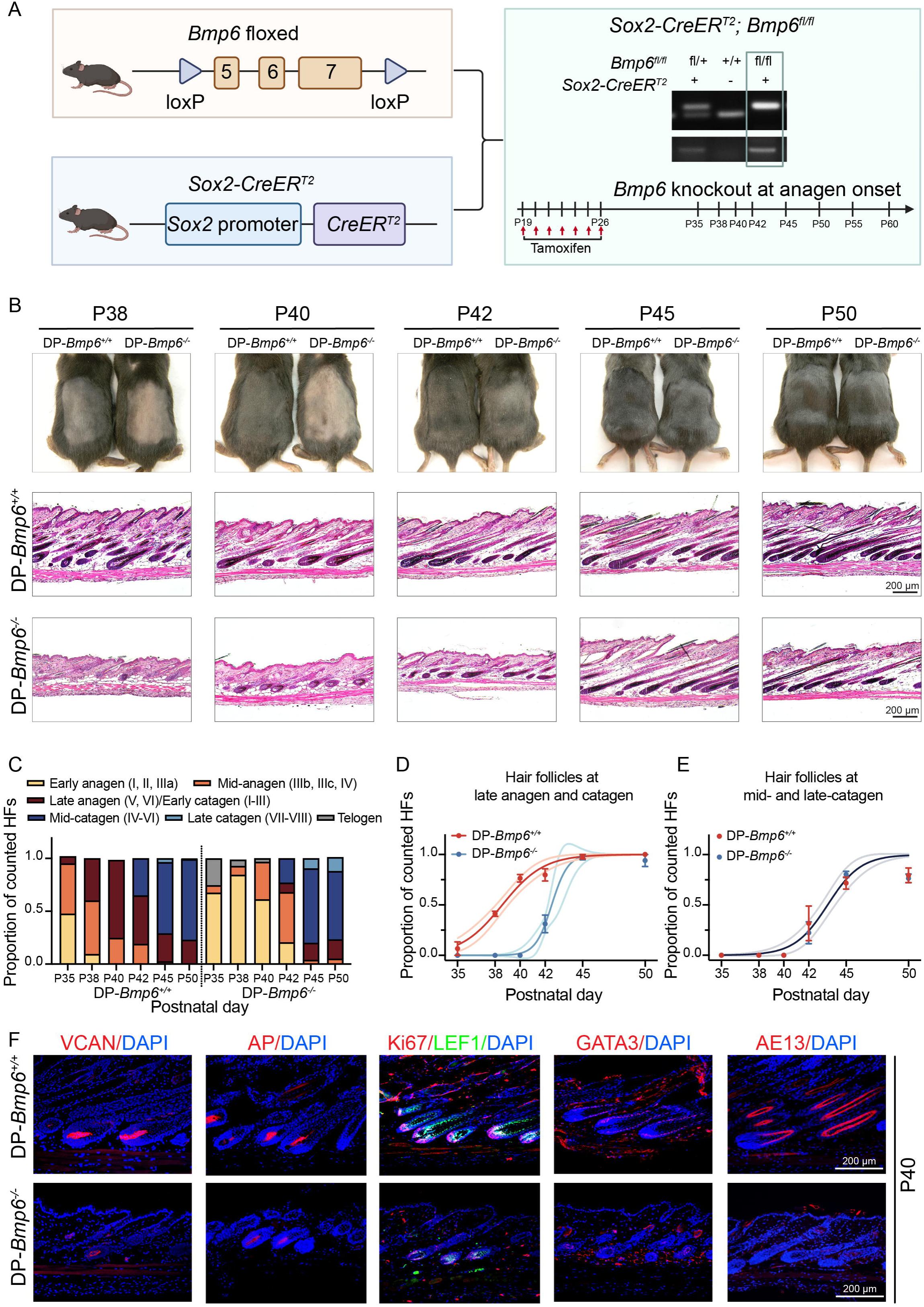
*Bmp6* deletion in *Sox2*-expressing DP cells delays anagen onset and disrupts anagen progression and follicular differentiation. **A.** Schematic of the conditional knockout model generated by crossing *Sox2-CreER^T^*^2^ and *Bmp6^fl/fl^*mice, with tamoxifen administration beginning at P19 to delete *Bmp6* in Sox2-expressing DP cells at anagen onset. The experimental timeline indicates tamoxifen administration and subsequent tissue-collection time points. **B.** Gross appearance and representative H&E-stained skin sections from *Sox2-CreER^T^*^2^; *Bmp6^fl/fl^*mice (DP-*Bmp6^-/-^*) and control littermates (*Sox2-CreER^T^*^2^; *Bmp6^+/+^*, DP-*Bmp6^+/+^*) at P38, P40, P42, P45, and P50. At least three mice per genotype were examined at each time point. Scale bars = 200 μm. **C.** Distribution of hair follicles across defined anagen, catagen, and telogen stages from P35 to P50 between DP-*Bmp6^+/+^* and DP-*Bmp6^-/-^*follicles. Five histological fields per mouse were quantified from three mice per genotype at each time point. Y-axis corresponds to the proportion of all counted hair follicles from 0 to 1. **D.** Non-linear curve fit of hair follicles in late anagen/early catagen, mid-catagen, and late catagen demonstrated a significant delay in anagen progression following *Bmp6* deletion in Sox2-positive cells (ΔT_50_ = 3.95 ± 0.66 days). Y-axis corresponds to the proportion of all counted hair follicles from 0 to 1, with data points corresponding to the mean per-mouse proportion of counted hair follicles in late anagen/early catagen, mid-catagen, and late catagen. **E**. Non-linear curve fit of hair follicles in mid- and late catagen showed no significant difference in catagen progression between DP-*Bmp6^+/+^* and DP-*Bmp6^-/-^* follicles (ΔT_50_ = −0.15 ± 1.45 days). Y-axis corresponds to the proportion of all counted hair follicles from 0 to 1, with data points corresponding to the mean per-mouse proportion of counted hair follicles in mid- and late catagen. **F.** Representative immunostaining for VCAN, AP, GATA3, LEF1/Ki67, and AE13 expression in DP-*Bmp6^+/+^* and DP-*Bmp6^-/-^*hair follicles at P40, when the phenotypic difference between groups was most pronounced. Scale bars = 200 μm.

At P40, when the phenotypic divergence was most pronounced, VCAN expression and AP activity were markedly reduced in the DP of follicles in DP-*Bmp6^-/-^*mice, consistent with attenuation of the mature DP phenotype (Fig. 2F). These DP defects were accompanied by diminished LEF1 expression in the hair matrix, markedly reduced GATA3 expression in the IRS, and near-absence of AE13-positive cortex keratins, indicating a profound disruption of follicular epithelial differentiation. Thus, the failure of mutant follicles to reach full anagen was accompanied by defective DP maturation and coordinated epithelial differentiation.

To understand whether *Bmp6* is also required after anagen has been established, we induced *Bmp6* deletion beginning at P28 (Fig. S1A). Follicles in DP-*Bmp6^-/-^* mice subsequently underwent accelerated regression, with progressive follicle shortening and bulb involution evident by P42–P45 (Fig. S1B). Quantitative staging demonstrated a shift toward more advanced catagen stages in mutant follicles, followed by earlier entry into telogen, with a markedly greater proportion of mutant follicles in telogen by P50 (Fig. S1C, D). As shown in Fig. S1E, VCAN, Ki67, and LEF1 expression was reduced in mutant hair follicles, with minimal changes in GATA3, AE13, and TCHH. Collectively, the two induction paradigms indicate that *Bmp6* in Sox2-expressing DP cells contributes to both anagen maturation and maintenance. We next asked whether BMP6 could restore the DP characteristics progressively lost during expansion of human DP fibroblasts *in vitro*.

### BMP6 restores culture-attenuated characteristics of human DP fibroblasts

Early-passage human DP fibroblasts are highly proliferative and express AP, a marker associated with hair-inductive competence^48^. Conventional monolayer expansion progressively erodes the DP phenotype, with P5–P6 cells exhibiting substantial reductions in AP activity, VCAN expression, and aggregative behavior^49^. Recombinant human BMP6 increased AP activity in both P3 and P5 monolayer cultures, restoring activity in P5 cells toward levels observed in untreated P3 cells (Fig. 3A, B). BMP6 also significantly increased cell expansion in both passages by day seven (Fig. 3C), accompanied by a higher proportion of Ki67-positive cells (Fig. 3D, E). Thus, exogenous BMP6 enhances both proliferative activity and AP expression in cultured human DP fibroblasts.

**Figure 3.**
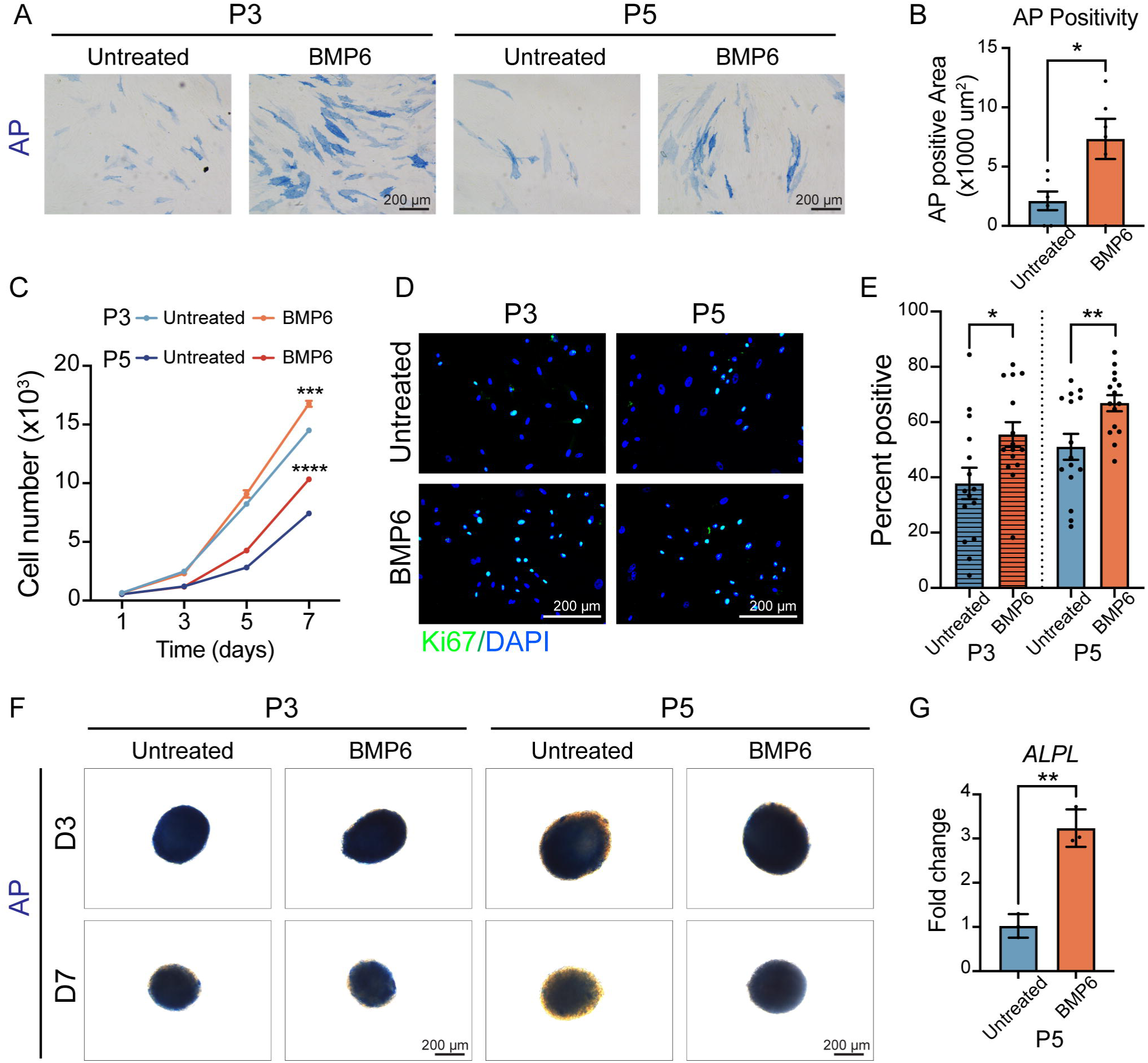
Recombinant BMP6 preserves DP-associated characteristics of human DP fibroblasts. **A.** Representative AP staining images of P3 and P5 DP fibroblasts cultured with or without recombinant BMP6 for three days. Scale bars = 200 μm. **B.** Quantification of AP-positive area in P5 DP fibroblasts cultured with or without BMP6 for three days. Each data point represents the AP-positive area measured in one field. Six fields per treatment condition were analyzed across three donor-derived DP fibroblast strains, and data are presented as mean ± SEM. **C.** Proliferation of P3 and P5 DP fibroblasts cultured with or without BMP6 across seven days. At day seven, BMP6 significantly increased cell number in both P3 cultures (14,496 ± 198.5 vs. 16,760 ± 254.6) and P5 cultures (7,423 ± 112.6 vs. 10,335 ± 174.7). Data are presented as mean ± SEM from four independent experiments. **D.** Representative Ki67 immunostaining of P3 and P5 DP fibroblasts cultured with or without BMP6 for three days. Scale bars = 200 μm. **E.** Quantification of Ki67-positive cells showing significantly increased Ki67 positivity following BMP6 treatment in both P3 (37.80% ± 5.65 vs. 55.50% ± 4.49) and P5 (51.05% ± 4.71 vs. 66.85% ± 2.92) DP fibroblasts. Each data point represents one field. 15 fields per treatment condition were analyzed across three donor-derived DP fibroblast strains. Data are presented as mean ± SEM. **F.** Representative AP staining images of P3 and P5 DP spheroids after three or seven days of culture with or without BMP6. Scale bars = 200 μm. **G.** RT-qPCR analysis of *ALPL* expression in P5 DP spheroids after three days of culture with or without BMP6. *ALPL* expression increased 2.21 ± 0.29-fold following BMP6 treatment. Data are presented as mean ± SEM from three independent experiments. * *p* < 0.05, ** *p* < 0.01, *** *p* < 0.001, **** *p* < 0.0001; n.s., not significant.

In 3D culture, AP activity declined with passage and culture duration. P3 spheroids retained substantial AP activity through day seven, whereas P5 spheroids exhibited reduced activity at day three and minimal staining by day seven (Fig. 3F). BMP6 had little additional effect in P3 spheroids but markedly increased AP activity in P5 spheroids at day seven, indicating that its restorative effect was most pronounced after culture-associated attenuation of DP characteristics. Consistent with this effect, *ALPL* expression was significantly increased in BMP6-treated P5 spheroids compared with untreated controls (Fig. 3G).

### BMP6 maintains and restores DP-associated characteristics across culture expansion

To define the role of endogenous BMP6 during progressive culture expansion, we used complementary loss- and gain-of-function approaches at passages representing different levels of retained DP competence. In early-passage P4 DP fibroblasts, where DP characteristics were still relatively retained, *BMP6* was depleted using doxycycline-inducible shRNA to test its role in phenotype maintenance. In P5 DP fibroblasts, where DP characteristics were attenuated, endogenous BMP6 was activated using CRISPRa to determine whether increasing BMP6 expression could restore the declining phenotype. BMP6 depletion and activation were validated at the transcript or protein levels (Fig. 4D, F; S2A, B).

**Figure 4.**
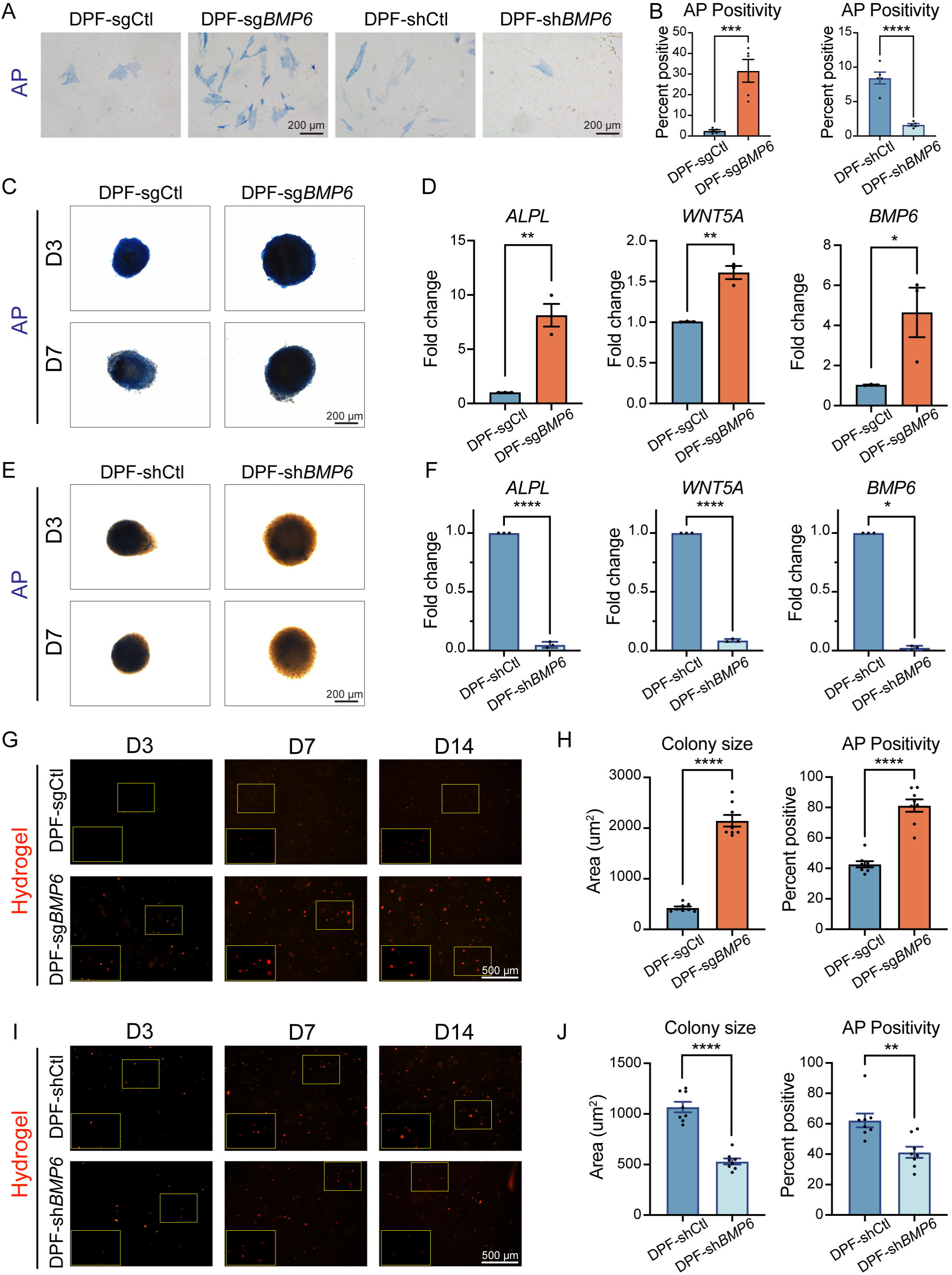
Endogenous BMP6 activation and silencing have opposing effects on DP-associated characteristics in human DP fibroblasts. **A.** Representative AP staining images of P5 CRISPRa control cells (DPF-sgCtl), P5 BMP6-activated cells (DPF-sg*BMP6*), P4 shRNA control cells (DPF-shCtl), and P4 BMP6-knockdown cells (DPF-sh*BMP6*). Scale bars = 200 μm. **B.** Quantification of AP-positive area as a percentage of total cell-covered area in monolayer cultures. AP-positive area was increased in DPF-sg*BMP6* compared with DPF-sgCtl (31.57% ± 5.51 vs. 2.56% ± 0.53) and decreased in DPF-sh*BMP6* compared with DPF-shCtl (1.63% ± 0.18 vs. 8.43% ± 0.86). Each data point represents the AP-positive area measured in one field. Five fields per group were analyzed across three independent experiments, and data are presented as mean ± SEM. **C.** Representative AP staining images of DPF-sg*BMP6* and DPF-sgCtl spheroids after three and seven days of culture. Scale bars = 200 μm. **D.** RT-qPCR analysis showing increased *ALPL*, *WNT5A*, and *BMP6* expression in DPF-sg*BMP6* spheroids. Data are presented as mean ± SEM from three donor-derived DP fibroblast strains per condition. **E.** Representative AP staining images of DPF-sh*BMP6* and DPF-shCtl spheroids after three and seven days of culture. Scale bars = 200 μm. **F.** RT-qPCR analysis showing decreased *ALPL*, *WNT5A*, and *BMP6* expression in DPF-sh*BMP6* spheroids. Data are presented as mean ± SEM from three donor-derived DP fibroblast strains per condition. **G.** Representative AP staining images of hydrogel-cultured DPF-sgCtl and DPF-sg*BMP6* at days three, seven, and 14. The yellow box at the lower left shows an enlarged view of the boxed region in the image. Scale bars = 500 μm. **H.** At day 14, DPF-sg*BMP6* colonies were larger than DPF-sgCtl colonies (2146 ± 115.6 vs. 425.6 ± 27.65 μm²) and showed greater AP positivity (81.25% ± 4.08 vs. 42.69% ± 2.07). For colony size, each data point represents the mean colony area within one field. For AP positivity, each data point represents the percentage of AP-positive colonies relative to the total number of colonies within the corresponding field. Eight randomly selected fields per group, obtained across three independent experiments, were analyzed. **I.** Representative AP staining images of hydrogel-cultured DPF-shCtl and DPF-sh*BMP6* at days three, seven, and 14. The yellow box at the lower left shows an enlarged view of the boxed region in the image. Scale bars = 500 μm. **J.** At day 14, DPF-sh*BMP6* colonies were smaller than DPF-shCtl colonies (530 ± 29.49 vs. 1068 ± 52.04 μm²) and showed lower AP positivity (41.25% ± 3.65 vs. 62.20% ± 4.56). For colony size, each data point represents the mean colony area within one field. For AP positivity, each data point represents the percentage of AP-positive colonies relative to the total number of colonies within the corresponding field. Eight randomly selected fields per group, obtained across three independent experiments, were analyzed. * p < 0.05, ** p < 0.01, *** p < 0.001, **** p < 0.0001; n.s., not significant.

CRISPRa-mediated *BMP6* activation significantly increased AP activity in monolayer P5 DP fibroblasts compared with non-targeting controls, reproducing the restorative effect of recombinant BMP6 (Fig. 4A, B). Conversely, BMP6 depletion substantially reduced AP positivity in P4 DP fibroblasts. In spheroid culture, AP staining was comparable between control and BMP6-activated groups at day three but declined markedly in control spheroids by day seven while remaining strongly retained following BMP6 activation (Fig. 4C). BMP6 activation was accompanied by significant increases in the DP-associated genes *ALPL* and *WNT5A*, together with increased *BMP6* expression (Fig. 4D). Conversely, BMP6 depletion markedly reduced AP activity in P4 spheroids at both day three and day seven (Fig. 4E), accompanied by profound reductions in *ALPL*, *WNT5A*, and *BMP6* expression (Fig. 4F).

We further examined whether these reciprocal effects were maintained in hydrogel culture. BMP6-activated DP fibroblasts progressively formed larger colonies than non-targeting controls and exhibited substantially greater AP positivity (Fig. 4G, H). Conversely, BMP6-depleted DP fibroblasts formed smaller colonies and showed reduced AP positivity compared with shRNA controls (Fig. 4I, J). Together, these reciprocal gain- and loss-of-function studies demonstrate that endogenous BMP6 contributes to maintenance of DP-associated characteristics, whereas increasing BMP6 expression enhances features lost during culture expansion across monolayer, spheroid, and hydrogel culture environments.

### BMP6 restricts proliferation and alters morphology in human epidermal KCs

In contrast to its proliferative effects on DP fibroblasts, BMP6 has previously been shown to suppress proliferation of primary human epidermal KCs^50^. Consistent with these reports, recombinant BMP6 reduced KC expansion over seven days in our cultures (Fig. 5A) and markedly decreased Ki67 positivity after 72 hours of treatment. Elevated extracellular calcium is known to suppress KC proliferation^51^, but it did not produce a comparable reduction in Ki67 positivity under the conditions examined (Fig. 5B, C). BMP6 treatment was associated with decreased expression of multiple cell-cycle regulatory proteins, including cyclin A2, cyclin B1, cyclin D3, cyclin E, CDK2, and CDK4. In contrast, CaCl_2_ treatment primarily reduced cyclin A2, with relatively limited changes in the other cell-cycle regulators examined (Fig. 5D).

**Figure 5.**
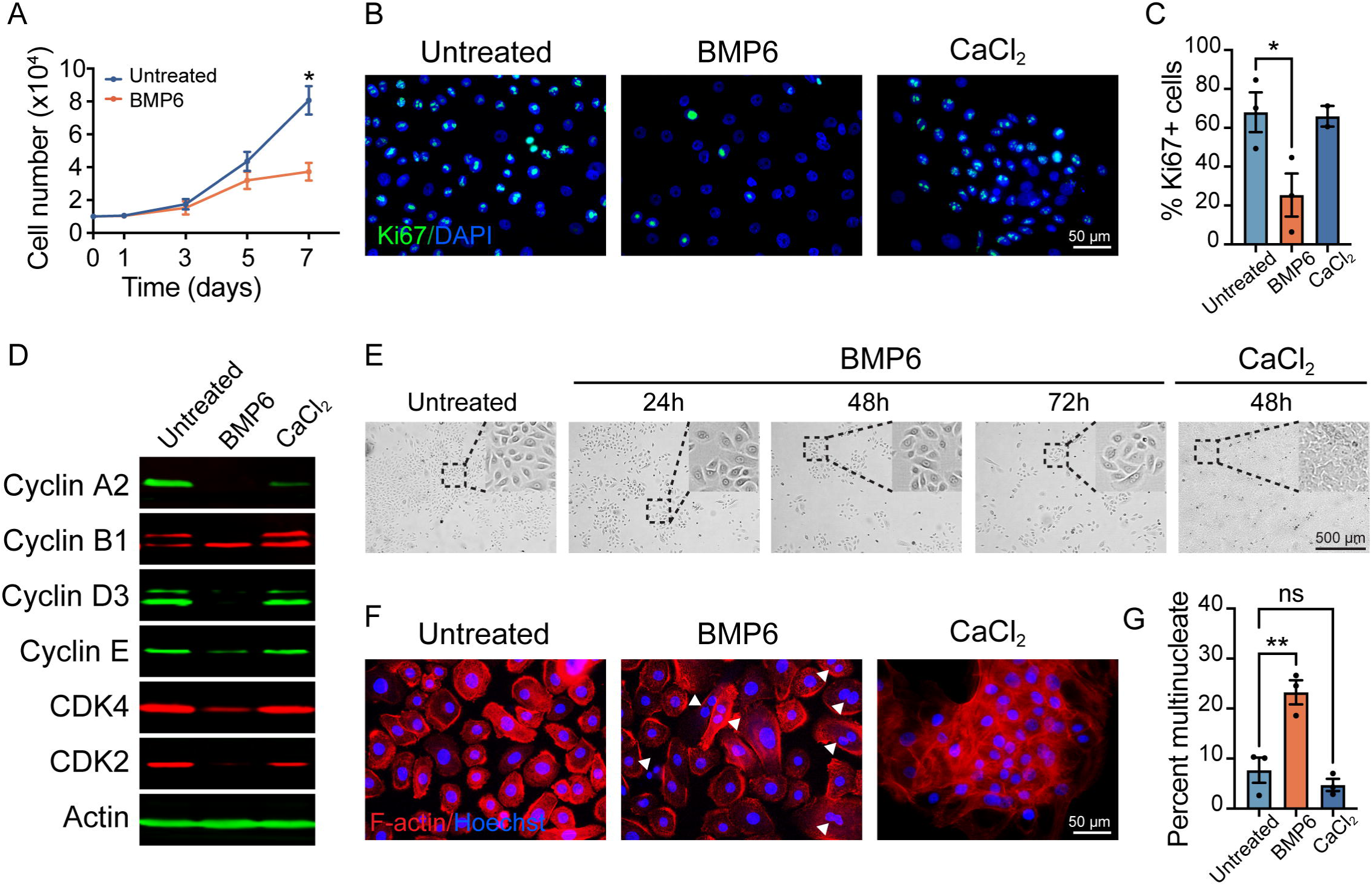
BMP6 restricts proliferation and alters morphology in human epidermal KCs. **A.** Proliferation of human KCs cultured with or without 200 ng/mL BMP6 over seven days. BMP6 significantly reduced cell number at day seven. Data are presented as mean ± SEM from three independent experiments. **B.** Representative Ki67 immunostaining of untreated KCs and KCs treated with 500 ng/mL BMP6 for 72 hours or 2 mM CaCl for 48 hours. Scale bars = 50 μm. **C.** Quantification of Ki67-positive cells showing reduced Ki67 positivity following BMP6 treatment, whereas CaCl treatment did not produce a comparable reduction. Each data point represents the average of five fields of view per sample, with samples derived from at least three donor-derived KC strains. **D.** Western blot analysis of cyclins A2, B1, D3, and E and CDK2 and CDK4 in untreated, BMP6-treated, and CaCl -treated KCs. Actin served as the loading control. Blots are representative of experiments performed using three donor-derived KC strains. **E.** Phase-contrast images of untreated KCs, KCs treated with BMP6 for 24, 48, or 72 hours, and KCs treated with CaCl for 48 hours, showing progressive morphological changes following BMP6 treatment. Scale bars = 500 μm. **F.** F-actin staining with rhodamine phalloidin and Hoechst nuclear counterstaining in untreated, BMP6-treated, and CaCl -treated KCs. Arrowheads indicate multinucleated cells observed following BMP6 treatment. Scale bars = 50 μm. **G.** Quantification of multinucleated cells showing a significant increase following BMP6 treatment (7.67% ± 2.53 vs. 23.25% ± 2.41), whereas CaCl treatment did not significantly alter multinucleation (7.67% ± 2.53 vs. 4.72% ± 1.22). Each data point represents the average of five fields of view per sample, with three donor-derived KC strains analyzed per condition. * *p* < 0.05, ** *p* < 0.01, *** *p* < 0.001, **** *p* < 0.0001; n.s., not significant.

The cellular morphology observed following the two treatments also differed. Calcium-treated KCs became flattened and densely packed, characteristic of conventional epidermal differentiation, whereas BMP6-treated cells retained a morphology more similar to untreated KCs despite reduced proliferation (Fig. 5E). BMP6 treatment was also associated with an increased frequency of multinucleated KCs, an effect not observed with calcium (Fig. 5F, G). Together, these observations confirm the growth-restrictive effect of BMP6 in human KCs and show that BMP6-treated cells exhibit morphological features that differ from those observed after calcium treatment. We therefore next examined the nature of the BMP6-induced KC differentiation response and whether it differed from conventional calcium-induced epidermal differentiation.

### BMP6 promotes follicular epithelial differentiation in human KCs

We compared the expression of epidermal and follicular differentiation markers in KCs treated with BMP6 or calcium. The suprabasal epidermal marker K10 remained undetectable following BMP6 treatment, whereas BMP6 induced expression of the hair cortex-associated keratin recognized by AE13, which was absent in untreated KCs (Fig. 6A, B). At the transcript level, BMP6 increased *TCHH* expression without inducing *KRT10*, whereas calcium increased *KRT10* but not *TCHH* (Fig. 6C). Because these experiments were performed in the absence of DP fibroblasts, these findings indicate that BMP6 can act directly on human epidermal KCs and promote acquisition of follicular epithelial characteristics distinct from calcium-induced epidermal differentiation.

**Figure 6.**
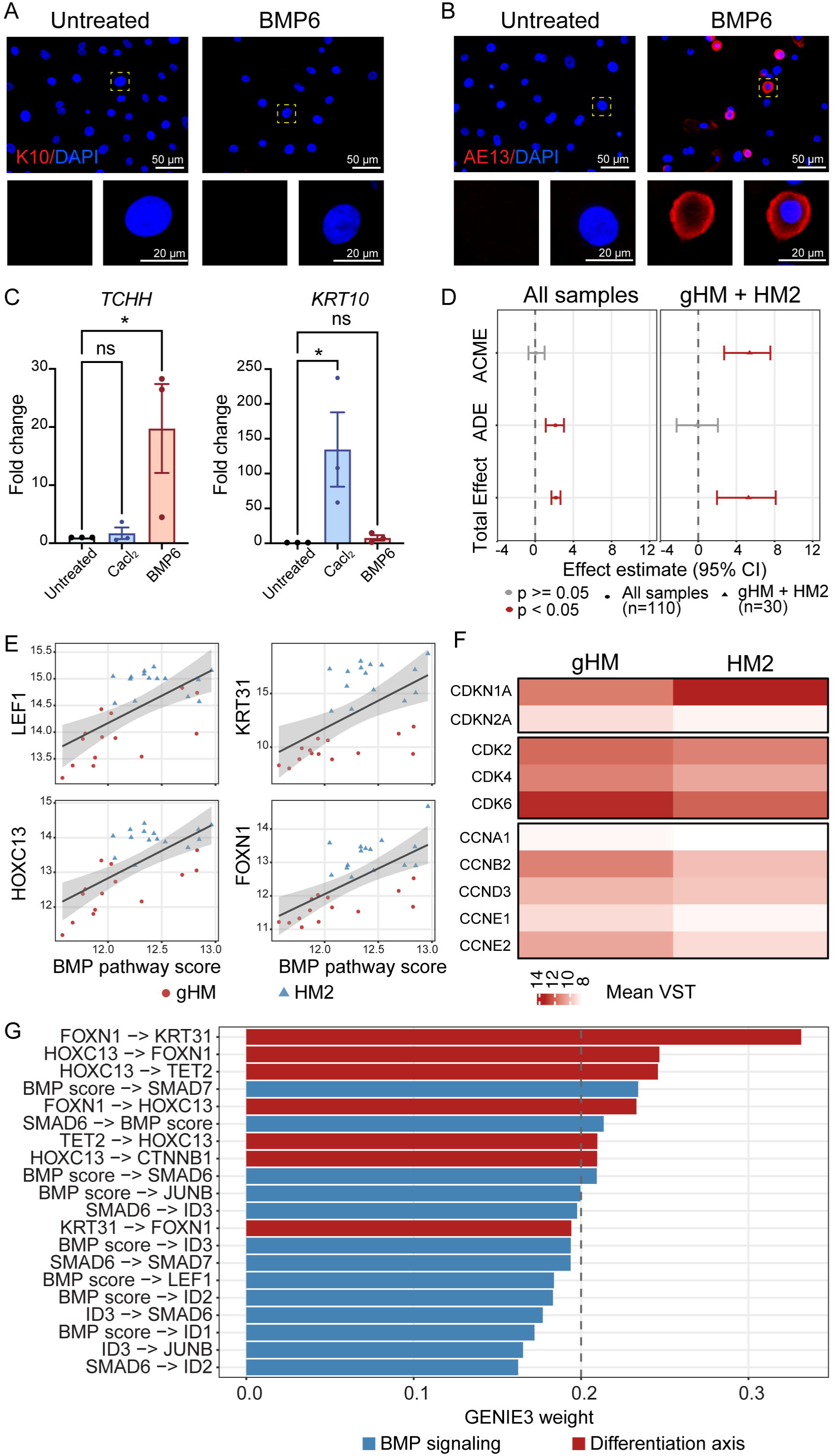
BMP6 promotes follicular epithelial differentiation *in vitro* and is associated with differentiation programs in human hair follicles. **A.** Immunostaining of untreated and BMP6-treated human epidermal KCs for the suprabasal epidermal differentiation marker K10. Scale bars = 50 μm and 20 μm. **B.** Immunostaining of untreated and BMP6-treated KCs for the follicular differentiation marker AE13. Scale bars = 50 μm and 20 μm. **C.** RT-qPCR analysis shows increased *TCHH* expression after BMP6 treatment (*p* < 0.05) and increased *KRT10* expression after CaCl treatment; the reciprocal comparisons were not significant. Data are presented as mean ± SEM from three donor-derived KC strains. **D.** Mediation analysis examining the association between the BMP pathway score and *KRT31* expression through *LEF1*. Across all follicular compartments, the indirect effect through *LEF1* was not significant, whereas the direct effect was significant. In the hair bulb compartments (gHM and HM2), the indirect effect was significant, whereas the direct effect was not significant; the estimated proportion mediated was 1.022 (95% CI 0.68–1.77). **E**. The BMP pathway score was positively correlated with *LEF1* (ρ = 0.62), *KRT31* (ρ = 0.61), *HOXC13* (ρ = 0.64), and *FOXN1* (ρ = 0.67) across gHM and HM2. **F**. Heatmap of cell cycle-associated genes in gHM and HM2 showing increased *CDKN1A* and reduced expression of multiple cell-cycle regulators, including *CDK2*, *CDK4*, *CDK6*, *CCNB2*, *CCND3*, and *CCNE2*, in HM2 relative to gHM. **G**. GENIE3 analysis across all 110 samples showing predicted relationships among BMP-responsive and follicular differentiation-associated genes. *FOXN1*→*KRT31* and *HOXC13*→*FOXN1* were among the highest-ranked differentiation-associated relationships, while the BMP pathway score was associated with *SMAD6*, *SMAD7*, *JUNB*, *ID* genes, and *LEF1*. * *p* < 0.05, ** *p* < 0.01, *** *p* < 0.001, **** *p* < 0.0001; n.s., not significant.

To determine whether this response was associated with epithelial differentiation in native human follicles, we analyzed the hair matrix and precortex compartments of GSE272333, which represent successive stages of matrix KC differentiation. A BMP pathway score derived from eight BMP-responsive genes was calculated for this analysis. Mediation analysis was used to examine the association between the BMP pathway score and *KRT31* expression through *LEF1* (Fig. 6D). Across all follicular compartments, the indirect effect through *LEF1* was not significant, whereas the direct effect was significant. Within gHM and HM2, however, the indirect effect was significant whereas the direct effect was not, consistent with an association of *LEF1* with the BMP pathway score–*KRT31* relationship in these hair bulb compartments. The BMP pathway score was also positively correlated with *LEF1* and with the differentiation-associated genes *KRT31, HOXC13,* and *FOXN1* across gHM and HM2 (Fig. 6E). Consistent with the transition toward differentiation, HM2 showed increased *CDKN1A* and reduced expression of multiple cell-cycle regulators compared with gHM (Fig. 6F), consistent with progression from proliferative matrix KCs toward precortex differentiation.

GENIE3 analysis further identified predicted relationships among BMP-responsive genes, *LEF1*, and follicular differentiation-associated genes (Fig. 6G). Among the higher-ranked relationships were *FOXN1*→*KRT31* and *HOXC13*→*FOXN1*, while the BMP pathway score was associated with *SMAD6, SMAD7, JUNB*, ID genes, and *LEF1*. Together, these compartment-resolved human follicle data associate the BMP pathway score with a growth-restricted, follicular differentiation-associated epithelial state and provide a native human follicle context consistent with the response observed in BMP6-treated KCs.

### BMP6 restores epithelial–mesenchymal organization and follicular differentiation in late-passage KC–DP organoids

We evaluated KC–DP organoid formation using recombinant BMP6 treatment and CRISPRa-mediated *BMP6* activation in P5 DP fibroblasts and shRNA-mediated *BMP6* depletion in P4 DP fibroblasts, followed by lentiviral *BMP6* overexpression in more extensively expanded P6 cells. P3 DP fibroblasts co-cultured with epidermal KCs develop hair peg-like structures through successive self-organization stages of aggregation, polarization, budding, and elongation (Fig. 7A). While P5 DP fibroblasts incompletely polarized and thus did not form advanced hair peg-like structures with KCs, addition of recombinant BMP6 to P5 KC–DP co-cultures partially restored polarization and initiated hair peg-like structure formation. CRISPRa-mediated activation of endogenous *BMP6* produced a more pronounced rescue, yielding polarized structures resembling those generated by P3 DP fibroblasts. Conversely, BMP6 depletion disrupted organization and prevented formation of comparable polarized hair peg-like structures (Fig. 7B). These organizational phenotypes were accompanied by corresponding changes in AP activity: P5 control organoids showed smaller and weaker AP-positive regions; recombinant BMP6 increased AP activity, CRISPRa-mediated BMP6 activation restored a large AP-positive DP compartment, and BMP6 depletion markedly reduced AP activity (Fig. 7C).

**Figure 7.**
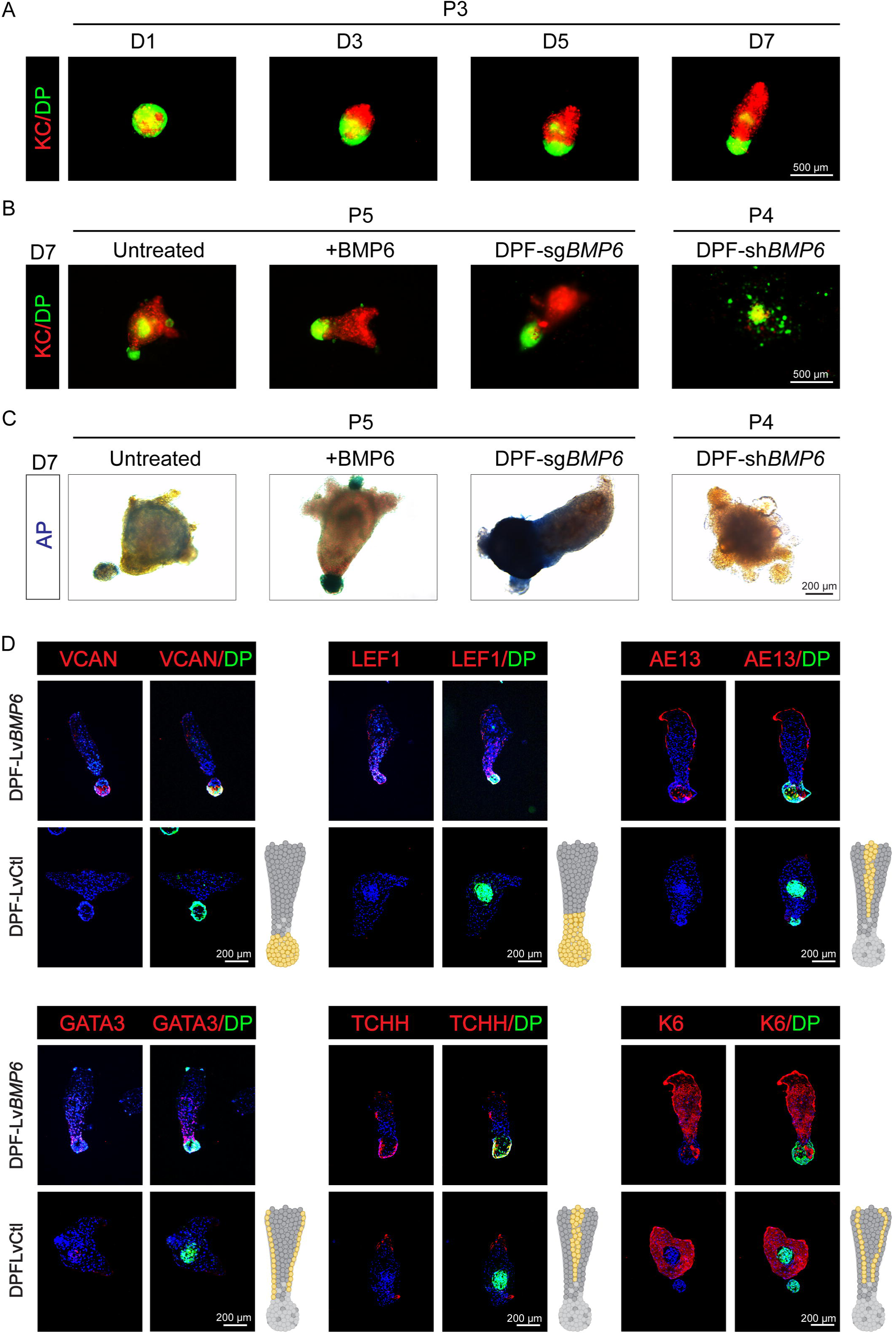
BMP6 enables late-passage human DP fibroblasts to form hair peg-like structures with KCs. **A.** Time course of hair peg-like structure formation from P3 DP fibroblasts across seven days, with DP fibroblasts labeled in green and KCs labeled in red. Scale bars = 500 μm. **B.** Representative day seven images of KC–DP composite spheroids generated with P5 non-transduced/untreated DP fibroblasts, P5 BMP6-treated DP fibroblasts, P5 DPF-sg*BMP6*, and P4 DPF-sh*BMP6*. DP fibroblasts are shown in green and KCs in red. P5 DP fibroblasts failed to form organized hair peg-like structures without BMP6, whereas BMP6 treatment or CRISPRa-mediated BMP6 activation restored hair peg-like organoid formation; BMP6 knockdown disrupted this process. Scale bars = 500 μm. **C.** Representative AP staining images of day seven KC–DP spheroid structures corresponding to B, showing increased AP activity in organoids formed with BMP6-treated or BMP6-activated DP fibroblasts and reduced AP activity after BMP6 knockdown. Scale bars = 500 μm. **D.** Immunostaining of sectioned day seven KC–DP structures generated with P6 DPF-LvCtl and P6 DPF-Lv*BMP6* for VCAN, LEF1, AE13, GATA3, TCHH, and K6 expression. BMP6 overexpression restored organized hair peg-like architecture and follicular marker expression in late-passage KC–DP composite organoids. Scale bars = 200 μm. Yellow regions in the gray hair follicle schematics indicate reported *in vivo* marker localization. Images are representative of three independent experiments.

To determine whether BMP6 could improve DP cell competence after more extensive culture expansion, P6 DPF-Lv*BMP6* and DPF-LvCtl cells were compared. BMP6 overexpression was confirmed in P6 DPF-Lv*BMP6* cells (Fig. S2C). DPF-Lv*BMP6* cells showed increased AP activity compared with DPF-LvCtl cells (Fig. S2D), reaching levels qualitatively comparable to those observed in recombinant BMP6-treated DP fibroblasts (Fig. 3A) and DPF-sg*BMP6* cells (Fig. 4A). Furthermore, P6 DPF-Lv*BMP6* cells supported substantially greater hair peg-like organization than control P6 DP fibroblasts, accompanied by restoration of VCAN expression within the DP compartment and LEF1 expression in adjacent epithelial cells (Fig. 7D). AE13- and GATA3-positive epithelial populations were spatially organized in patterns consistent with hair cortex- and IRS-associated differentiation, respectively, while TCHH and K6 further demonstrated compartmentalized follicular epithelial differentiation.

Together, recombinant BMP6 treatment and two independent gain-of-function approaches enhanced the organizational competence of culture-expanded DP fibroblasts, whereas BMP6 depletion produced the opposite phenotype. These findings support the idea that BMP6 contributes to DP-associated competence and enables mixed KC–DP spheroids containing late-passage DP fibroblasts to develop spatially patterned follicular epithelial structures.

### BMP6 promotes anagen progression and restores the hair follicle-generating capacity of human DP fibroblasts *in vivo*

Having shown that loss of DP-derived *Bmp6* impairs anagen progression and maintenance, we next asked whether exogenous BMP6 could promote anagen entry and progression *in vivo*. B6 mice with telogen-stage dorsal hair follicles were treated topically with recombinant BMP6 or vehicle. After 10 days, BMP6-treated areas exhibited prominent skin darkening and hair shaft emergence, whereas matched control areas showed little or no regrowth (Fig. 8A, B). Histological analysis confirmed accelerated anagen progression, with BMP6-treated follicles exhibiting enlarged, deeply positioned bulbs, increased matrix pigmentation, and greater skin thickness, while control follicles largely exhibited characteristics of early anagen (Fig. 8C). Hair bulb diameter was increased following BMP6 treatment (Fig. 8D). This transition was accompanied by robust nuclear LEF1 expression and increased Ki67 staining in matrix KCs, consistent with anagen induction (Fig. 8E). Thus, topical BMP6 accelerated anagen entry and subsequent follicular progression relative to vehicle-treated skin.

**Figure 8.**
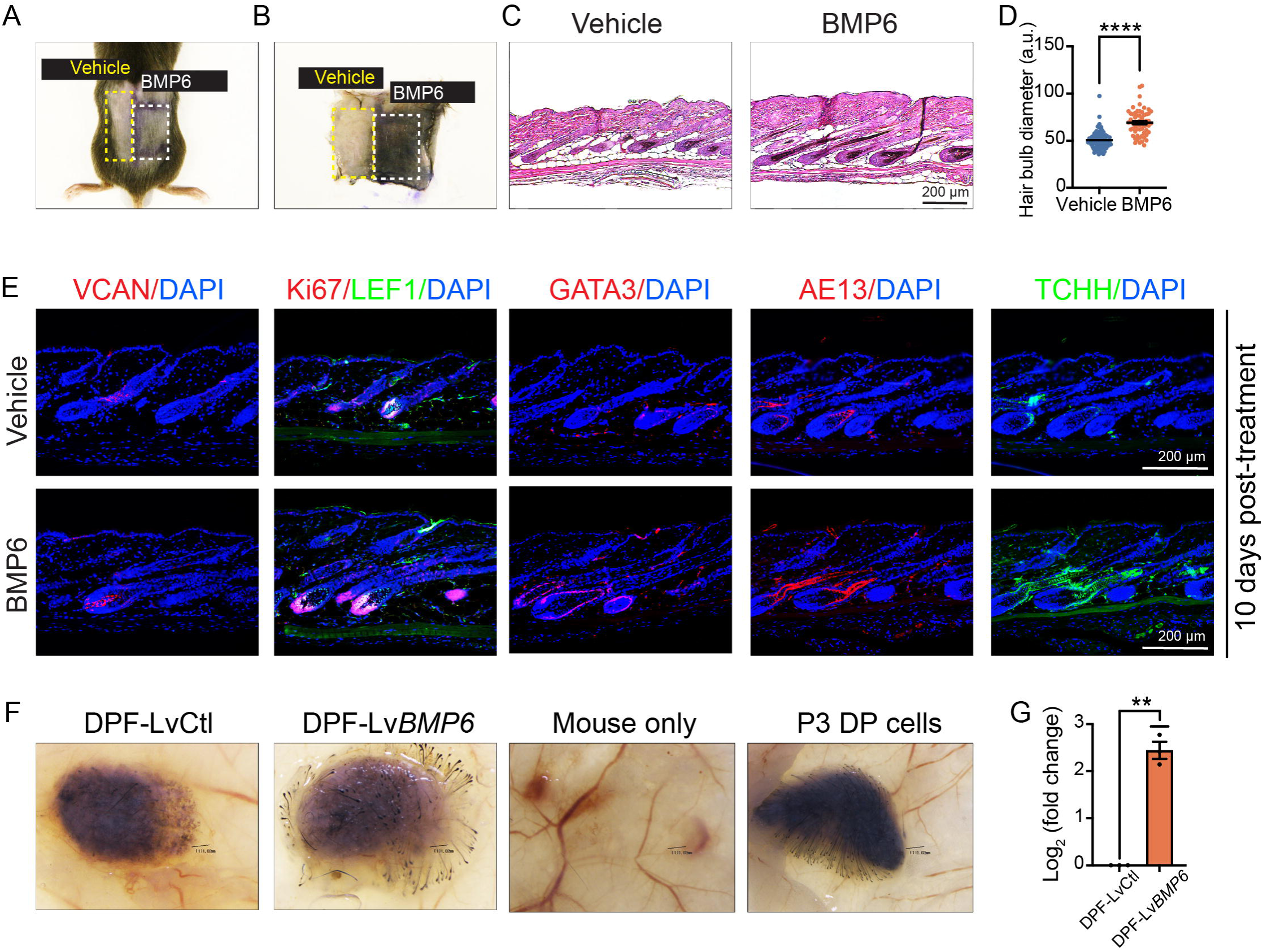
BMP6 promotes anagen-associated follicular activity and restores the hair-inductive capacity of late-passage human DP fibroblasts *in vivo*. A. Gross appearance of dorsal skin after topical treatment with 1 μg/cm^2^ BMP6 or vehicle for 10 days. **B.** Dermal-side view of BMP6-treated and vehicle-treated dorsal skin showing increased follicular pigmentation associated with anagen progression following BMP6 treatment. **C.** Histological morphology of BMP6-treated and vehicle-treated dorsal skin after 10 days of treatment, showing larger and more developed hair follicles following BMP6 treatment. Scale bars = 200 μm. Images are from one mouse showing paired vehicle-treated and BMP6-treated dorsal skin. **D.** Quantification of hair bulb diameter as a measure of anagen progression. BMP6 treatment significantly increased hair bulb diameter (50.72 ± 0.96 vs. 69.18 ± 1.77 a.u.). Hair bulb diameter was measured in follicles from five randomly selected skin fields per treatment condition across three mice, yielding a total of 92 vehicle-treated follicles and 60 BMP6-treated follicles. Individual follicle measurements were pooled by treatment condition for analysis. Data are presented as mean ± SEM. **E.** Representative immunostaining images of BMP6-treated and vehicle-treated dorsal skin for VCAN, GATA3, LEF1/Ki67, AE13, and TCHH, showing increased expression of anagen- and follicular differentiation-associated markers following BMP6 treatment. Scale bars = 200 μm. **F.** Representative dissection microscopy images of patch assays 28 days after intradermal injection of P7 control DPF-LvCtl or BMP6-overexpressing DPF-Lv*BMP6*, with mouse-only and P3 DP fibroblast groups included as negative and positive controls, respectively. BMP6 overexpression restored hair-forming activity in late-passage DP fibroblasts. **G.** Quantification of hair generation induced by DPF-LvCtl (14 ± 5.29 hairs) and DPF-Lv*BMP6* (83 ± 40.95 hairs). Within-mouse log fold change in hair regeneration for DPF-Lv*BMP6* relative to the contralateral DPF-LvCtl control was 2.44 ± 0.18 and was significantly greater than 0. Quantification was performed on three paired mice (n = 3 pairs of contralateral patches). * *p* < 0.05, ** *p* < 0.01, *** *p* < 0.001, **** *p* < 0.0001; n.s., not significant.

We next asked whether BMP6 could enhance the hair follicle-generating capacity of culture-expanded human DP fibroblasts. In an *in vivo* patch regeneration assay, late-passage P7 human DP fibroblasts were combined with neonatal C57BL/6 murine epidermal and dermal cells and injected intradermally into nude mice (Fig. 8F). P7 DPF-LvCtl generated an average of 14 hair follicles per patch, whereas passage-matched P7 DPF-Lv*BMP6* generated an average of 83 follicles per patch (Fig. 8G). Early-passage P3 DP fibroblasts retained appreciable follicle-generating activity and served as a positive reference for DP competence, whereas neonatal murine epidermal and dermal cells injected without human DP fibroblasts failed to generate follicles. These controls demonstrate that follicle generation depended on the inductive activity of human DP fibroblasts and that BMP6 overexpression substantially restored their hair follicle-generating capacity after prolonged culture expansion.

## Discussion

This study identifies BMP6 as a compartment-dependent coordinator of epithelial–mesenchymal interactions during hair growth. The follicle must preserve an active and instructive DP niche while simultaneously directing epithelial progenitors to withdraw from proliferation and differentiate into specialized follicular lineages. Our findings suggest that BMP6 helps coordinate these opposing requirements. In DP fibroblasts, BMP6 supports the specialized mesenchymal state and preserves epithelial-instructive competence. In KCs, the same signal restricts proliferation and promotes follicular rather than conventional epidermal differentiation. These complementary activities link DP maintenance with epithelial lineage formation, anagen onset, progression, and maintenance, and regenerative follicle function.

Rendl and colleagues established that BMP receptor signaling is required for murine DP identity and hair-inductive function and showed that exogenous BMP6 preserves AP activity and selected DP signature genes during culture^11^. The present findings extend this work by supporting an endogenous autocrine role for BMP6 in culture-expanded human DP fibroblasts. Complementary gain- and loss-of-function experiments in monolayer and spheroid cultures indicate that BMP6 is not merely an external enhancer of DP markers but contributes to maintenance of the AP-positive DP state. Importantly, BMP6 also supported DP fibroblast expansion, suggesting that it can partially uncouple proliferation from the progressive loss of specialized identity that normally accompanies monolayer culture. This property addresses a central problem in hair follicle engineering: DP fibroblasts must be expanded to obtain sufficient cell numbers, yet expansion diminishes the inductive competence for which the cells are required. The stronger restorative effect after passage-associated decline further suggests that BMP6 can re-establish elements of DP identity rather than simply reinforce an intact early-passage phenotype. Consistent with this interpretation, BMP6-overexpressing late-passage DP fibroblasts restored epithelial–mesenchymal organization in composite organoids and supported spatially organized matrix-, IRS-, and cortex-associated differentiation. Thus, evidence progressing from monolayer cultures to spheroids and composite organoids supports BMP6 as a niche-stabilizing signal that maintains an expandable and epithelial-instructive DP population.

Although multiple BMP ligands, including BMP2 and BMP4, contribute to hair follicle growth and differentiation^10,12,14^, BMP6 may possess properties that favor its activity within the DP–epithelial niche. BMP6 differs from BMP2/4 in receptor utilization in some cellular contexts and shows relative resistance to Noggin^20,52^, potentially allowing signaling to persist within the Noggin-rich follicular environment^16^. Its broader mesenchymal distribution places BMP6 within both the inductive DP and the surrounding DS, positioning it to influence DP function and epithelial–mesenchymal interactions across the follicular mesenchyme^46^. These properties may contribute to the prominent effects observed here, although direct comparisons with BMP2 and BMP4 will be required to establish ligand-specific functions in DP fibroblasts and KCs.

A central finding of this study is that BMP6 promotes a follicular differentiation response in epidermal KCs distinct from the conventional epidermal program induced by calcium^53^. BMP6 induced cortex-associated keratin expression without inducing K10, indicating that its effects extend beyond growth restriction to the acquisition of follicular epithelial characteristics. Although both BMP6 and calcium suppress KC proliferation, they engage different upstream signaling systems. Calcium-induced KC differentiation is mediated through calcium-sensing and intracellular calcium-dependent signaling, whereas BMP6 acts through BMP receptors and canonical SMAD signaling. These distinct signaling inputs may contribute to the different differentiation responses observed here, although the present study does not establish that they are responsible for this divergence.

This lineage bias may reflect the ability of BMP signaling to engage latent follicular competence in KCs and cooperate with transcriptional regulators of hair-shaft differentiation. Compartment-resolved human follicle transcriptomes showed that the BMP pathway score was positively associated with *LEF1* and with follicular differentiation-associated genes including *HOXC13, FOXN1,* and *KRT31*^54–58^. These relationships support the physiological relevance of the cultured KC phenotype by placing it within the molecular organization of follicular differentiation in intact human follicles, although the transcriptomic analyses do not establish a causal signaling hierarchy. The mechanism by which BMP6 selects this follicular differentiation program remains unresolved and warrants further investigation.

The opposite proliferative responses of DP fibroblasts and KCs are therefore biologically complementary rather than contradictory. Hair growth requires maintenance of an active, instructive DP niche while epithelial progenitors progressively exit the cell cycle and differentiate. BMP6 appears to support these distinct requirements simultaneously, preserving the proliferative and instructive DP compartment while restricting proliferation and promoting differentiation in epithelial cells. The organoid findings provide a functional demonstration of this coordination: restoration of the DP phenotype was accompanied by epithelial organization and differentiation within the composite organoids. BMP6 should therefore be viewed not as a uniformly pro- or antiproliferative factor, but as a context-dependent signal that aligns distinct cellular behaviors within the follicular unit.

The conditional mouse experiments place this compartment-specific model within the physiology of the hair cycle. Loss of *Bmp6* in Sox2-positive DP cells disrupted the mature DP phenotype and impaired matrix, IRS, and cortex differentiation. Anagen onset was delayed, subsequent progression to mature anagen was impaired, and mutant follicles entered regression without achieving the advanced anagen state observed in controls. This sequence supports the interpretation that *Bmp6* deficiency compromises coordinated DP–epithelial differentiation together with normal anagen onset and progression, rather than causing an isolated defect in hair-cycle timing. Deletion after follicles had already entered anagen further demonstrated that DP-derived BMP6 remains necessary during active growth. Anagen maintenance therefore appears to require continuous epithelial–mesenchymal coordination, not only transient signaling at follicle activation. Localized topical BMP6 provided complementary gain-of-function evidence by promoting anagen onset and subsequent growth of resting follicles. Together, the genetic loss- and pharmacological gain-of-function findings indicate that DP-derived BMP6 contributes to anagen onset, progression, and maintenance, whereas exogenous BMP6 can promote activation and growth of resting follicles.

Beyond regulation of existing follicles, BMP6 also addresses a distinct challenge in d*e novo* follicle regeneration. Restoration of AP, VCAN, or organoid morphology is biologically meaningful only if it corresponds to recovery of the defining function of DP fibroblasts—the ability to induce new follicle formation. The patch assay demonstrated that BMP6-overexpressing late-passage human DP fibroblasts exhibited substantial follicle-generating activity relative to passage-matched controls. Because neonatal murine skin cells alone did not generate follicles, this activity depended on inductive signals supplied by the human DP fibroblasts. The patch assay therefore provides a stringent functional endpoint beyond marker restoration and demonstrates increased trichogenic competence after culture expansion. Topical BMP6 and BMP6-modified DP fibroblasts therefore represent distinct regenerative applications: one promotes anagen onset in existing follicles, whereas the other restores the cellular competence required for *de novo* follicle generation. Together with the organoid findings, these results connect restoration of DP-associated characteristics with improved epithelial– mesenchymal organization and increased follicle-generating capacity.

These findings also define several important questions. The mechanisms by which BMP6 preserves DP competence while promoting follicular differentiation in KCs remain unresolved, including whether these compartment-specific responses reflect differences in receptor utilization, SMAD signaling, or lineage-specific transcriptional context. Direct comparisons with other follicular BMP ligands will be needed to determine whether the effects observed here are specific to BMP6. In addition, longer-term studies will be required to determine whether BMP6-restored DP fibroblasts support durable follicle maturation and cycling and whether localized BMP6 delivery can be optimized for regenerative application.

In conclusion, BMP6 functions not simply as a general hair-growth factor but as a context-dependent regulator of the distinct requirements of the follicular mesenchyme and epithelium. BMP6 supports an expandable and instructive DP niche while restricting epithelial proliferation and promoting follicular differentiation. This coordinated activity is associated with anagen onset, progression, and maintenance and can be harnessed either to promote anagen onset and growth in resting follicles or to enhance the *de novo* follicle-generating capacity of culture-expanded human DP fibroblasts. Together, these findings identify BMP6-dependent epithelial– mesenchymal coordination as an important component of follicular growth and regeneration and provide a framework for its application in hair follicle regenerative strategies.

## Acknowledgments

This work was supported by the National Institutes of Health (R01AR077238). The content is solely the responsibility of the authors and does not necessarily represent the official views of the National Institutes of Health.

## Author contributions

Conceptualization: TA, YZ; Methodology: LS, LZ, CD, YZ, DS, TA, JH; Investigation: LS, LZ, YC; Data analysis: LS, LZ, CD, YZ; Writing: LS, YZ; Review & Editing: LS, TA, DS, YZ; Figure preparation: LS, LZ, JW, YZ; Funding acquisition: YZ; Resources: DS, JB, SM, YZ; Supervision: YZ. All authors reviewed and approved the final manuscript. All authors agree to be accountable for all aspects of the work in ensuring that questions related to the accuracy or integrity of any part of the work are appropriately investigated and resolved and declare confidence in the integrity of the contributions of their coauthors.

## Data availability

The publicly available RNA-sequencing datasets analyzed in this study are available from the Gene Expression Omnibus under accession numbers GSE272333, GSE109256, GSE300176, GSE235480, and GSE136996. Other data supporting the findings of this study are available from the corresponding author upon reasonable request.

