## Supplemental Figures for "BMP6 coordinates dermal papilla and epithelial cell states for hair follicle engineering and regeneration"

**Table S1. Primary cell strains**

| Cell type | Age | Sex | Ethnicity | Site |
| --- | --- | --- | --- | --- |
| KCs | 14 | F | Caucasian | Breast |
| KCs | 17 | F | Caucasian | Breast |
| KCs | 15 | F | Black | Breast |
| KCs | 13 | F | Caucasian | Breast |
| DP fibroblasts | 67 | F | Caucasian | Temple |
| DP fibroblasts | 46 | F | Caucasian | Temple |
| DP fibroblasts | 50 | F | Caucasian | Temple |
| DP fibroblasts | 36 | M | Caucasian | Scalp |
| DP fibroblasts | 32 | M | Caucasian | Scalp |

| Name | Sequence | Genomic target, GRCh38 | Target strand | Position relative to transcriptional start site (7,726,099) |
| --- | --- | --- | --- | --- |
| sgBMP6-1 | GGACAGGCTGGCAGCCAATG | chr6:7,726,020–7,726,039 | - | −79 to −60 bp |
| sgBMP6-2 | GAGGCGACGCCTCTCGGGAG | chr6:7,725,944–7,725,963 | + | −155 to −136 bp |
| sgBMP6-3 | GCCGCCCCGCGGTAACCGAC | chr6:7,726,036–7,726,055 | - | −63 to −44 bp |
| Control sgRNA | GTGTAGTTCGACCATTCGTG |  |  |  |

**Table S2. BMP6 sgRNA and control sgRNA sequences**

| Species | Gene | Forward (5’-3’) | Reverse (5’-3’) |
| --- | --- | --- | --- |
| Human | *ALPL* | CGACGTCTTCACATTTGGTG | AGACTGCGCCTGGTAGTTGT |
| Human | *BMP6* | AACCTGGTGGAGTACGACAA | CGGGTGTCCAACAAAAATAG |
| Human | *WNT5A* | TAGCAGCATCAGTCCACAAA | CAAAACACGGCATCTCTCTT |
| Human | *TCHH* | CCTGAAGAACCTCCTTGAAAGGG | ACACGCCCATTACTGTCAAGATC |
| Human | *KRT10* | CCTGCTTCAGATCGACAATGCC | ATCTCCAGGTCAGCCTTGGTCA |
| Human | *ACTB* | GGACTTCGAAGAGATGG | AGCACTGTGTTGGCGTACAG |
| Human | *GAPDH* | GAGTCAACGGATTTGGTCGT | TTGATTTTGGAGGGATCTCG |
| Murine | *Bmp6* | AACCTGGTGGAGTACGACAA | CGGGTGTCCAACAAAAATAG |
| Murine | *Ctnnb1* | ATTGGCCTGTAGAGTTGCTG | AACAAGCAAGGCTAGGGTTT |
| Murine | *Actb* | AAGAGCTATGAGCTGCCTGA | TACGGATGTCAACGTCACAC |

**Table S3. RT-qPCR primer pairs**

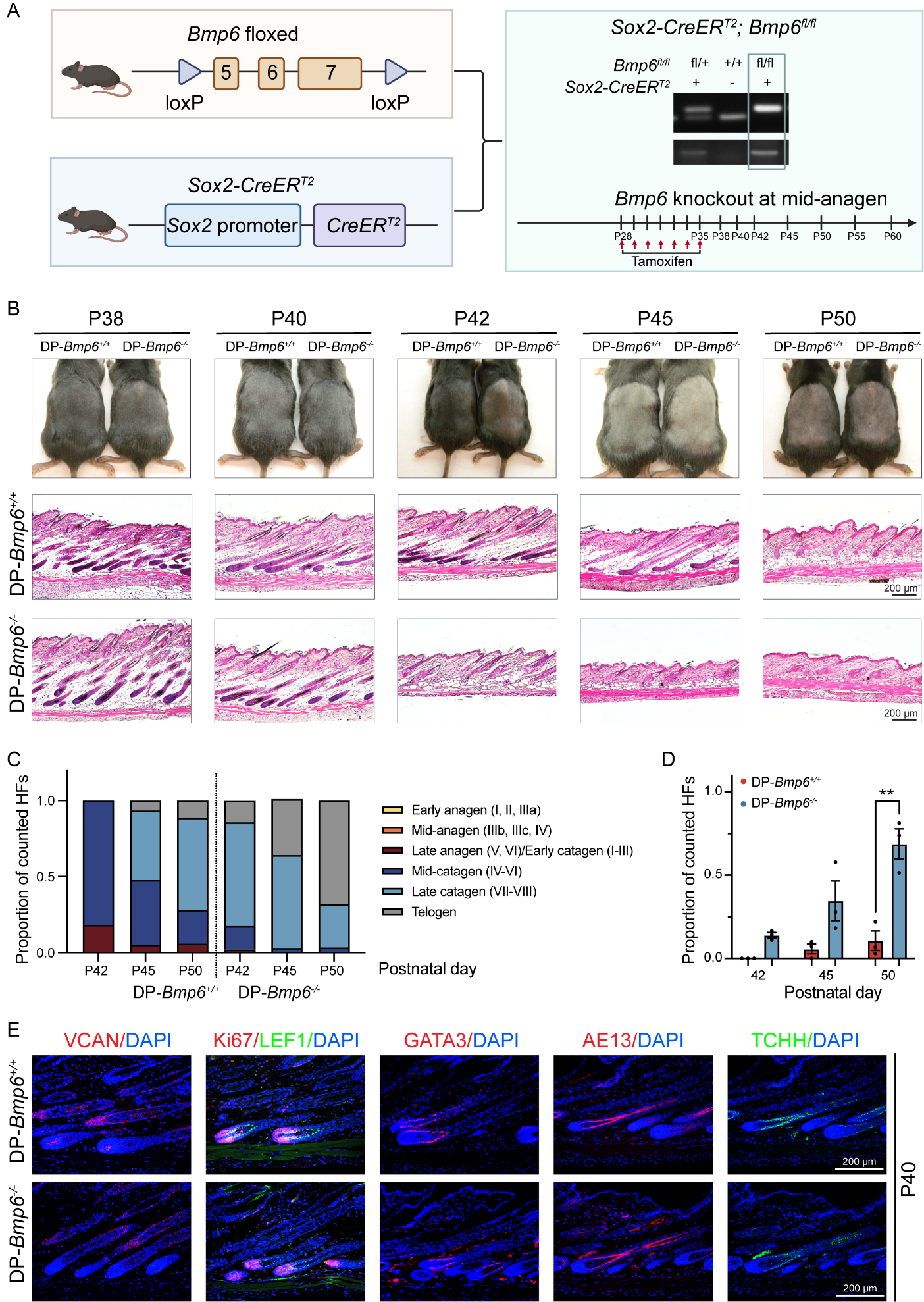

**Figure S1. *B*mp6 deletion in Sox2-positive DP cells during established anagen accelerates follicular regression.** **A.** Schematic of the Sox2-CreER^T2^; Bmp6*^fl/fl^* conditional knockout model, representative genotyping, and experimental timeline showing tamoxifen administration beginning at P28 to delete Bmp6 during established anagen. **B.** Gross appearance and representative H&E-stained dorsal skin from *Sox2-CreER^T2^*; *Bmp6^fl/fl^* mice (DP-Bmp6^-/-^) and control littermates (*Sox2-CreER^T2^*; *Bmp6^+/+^*, DP-Bmp6^+/+^) at P38, P40, P42, P45, and P50. DP-Bmp6^-/-^ follicles exhibit earlier regression than control follicles. Scale bars = 200 μm. **C.** Distribution of hair follicles across anagen, catagen, and telogen stages at P42, P45, and P50 following Bmp6 deletion in Sox2-positive cells initiated at P28. Five histological fields were quantified per mouse from three mice per genotype at each time point. The y-axis shows the proportion of all counted hair follicles on a scale from 0 to 1. **D.** Quantification of the proportion of follicles in telogen showing accelerated regression in DP-Bmp6^-/-^ mice, with a significant difference at P50 (*p* < 0.01). The y-axis shows the proportion of all counted hair follicles on a scale from 0 to 1; data points represent the mean per-mouse proportion of counted hair follicles at the designated time points. **E.** Representative immunostaining at P40 of DP-Bmp6^+/+^ and DP-Bmp6^-/-^ hair follicles for VCAN, Ki67/LEF1, GATA3, AE13, and TCHH expression. Scale bars = 200 μm. * *p* < 0.05, ** *p* < 0.01, *** *p* < 0.001, **** *p* < 0.0001; n.s., not significant.

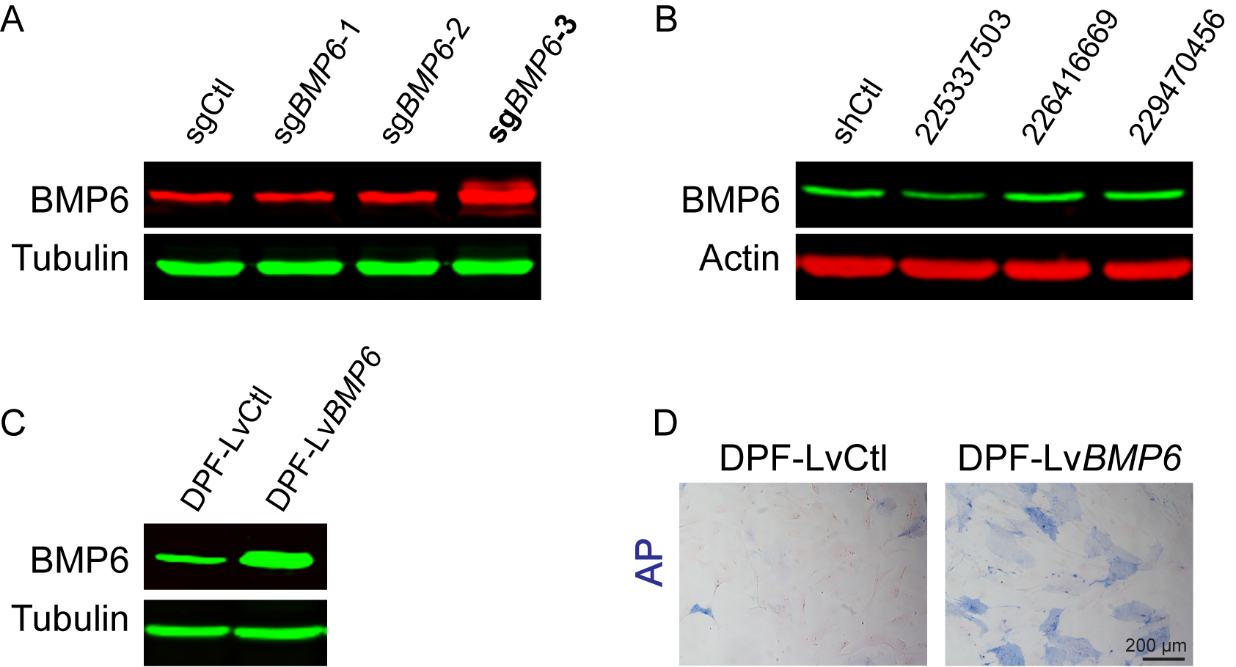

**Figure S2. Validation of BMP6 activation, knockdown, and overexpression in human DP fibroblasts. A.** Western blot analysis of BMP6 expression in P5 DP fibroblasts following transduction with three *BMP6*-targeting sgRNAs; sg*BMP6*-3 produced the greatest BMP6 activation and was selected for subsequent experiments. **B.** Western blot analysis of BMP6 expression in P4 DP fibroblasts following transduction with three *BMP6*-targeting shRNAs (V3SH7669-229470456, V3SH7669-226416669, and V3SH7669-225337503); V3SH7669-225337503 produced the greatest BMP6 knockdown and was selected for subsequent experiments. **C.** Western blot analysis confirming BMP6 overexpression following direct lentiviral transduction in P6 DPF‑LvCtl and DPF‑Lv*BMP6*. **D.** AP activity in P6 DPF-LvCtl and DPF-LvBMP6 following direct lentiviral BMP6 overexpression. Scale bar = 200 μm.
